# Characterisation of type 2 diabetes progression with regulatory networks

**DOI:** 10.1101/526954

**Authors:** María Barrera, Marcia Hiriart, Germinal Cocho, Carlos Villarreal

## Abstract

Type 2 diabetes develops due to beta cell exhaustion with accompanying decrease in insulin secretion, leading to hyperglycemia and eventual damage of nerve, kidney, and eye tissues. It is usually preceded by metabolic alterations related to insulin signaling, inflammatory pathways, or intracellular glucose processing, encompassed as metabolic syndrome. We propose a regulatory network for components of pancreatic-beta cells playing an essential role in the disease. The network interactions are expressed as continuous fuzzy logic propositions. The dynamical modeling of the network allows to portray the disease progression as a transit between steady states associated to health, metabolic syndrome, and diabetes, each state defined by specific expression patterns of the network components. Transitions between equilibrium states are due to altered expression or functional exhaustion associated to modifications of characteristic decay rates of cellular components. This approach let us identify functional modules that may eventually drive the transit from health to diabetes. The analysis reveals that underexpression of protein kinase B and X-box binding protein 1, with concomitant overexpression of lipopolysaccharides and thioredoxin interacting protein are key factors in the transition from health to disease.

## Introduction

Type 2 diabetes (T2D) is a significant public health problem around the world^1^ associated to dysregulation of carbohydrate, lipid, and protein metabolism^2^. T2D is characterized by a steady decline in insulin secretion from beta cells, resulting in elevated glucose levels that cause long-term complications in patients leading to eye, nerve, and kidney dysfunction. It is well documented that beta cells may undergo exhaustion by excessive insulin secretion caused by unsustainable demands^2^. As a consequence, some cells secrete less hormones while others may suffer premature programmed cell death, naturally resulting in a reduction of circulating insulin. T2D is usually preceded by metabolic syndrome (MS), constituted by a group of signs such as central obesity, dyslipidemia, glucose intolerance, hypertension, and insulin resistance, that increases the risk of developing T2D, as well as cardiovascular diseases and some types of cancer^3–5^. Furthermore, components of the immune response are also altered in obesity and T2D, including increased levels of inflammatory cytokines like interleukin 1 *β* (IL1*β*), interleukin 6 (IL6), and tumor necrosis factor (TNF), originating changes in the number and activation state of different leucocyte populations, increased apoptosis, and tissue fibrosis^6–8^. Other studies suggest an association of genes related to MS with molecular and physiological components involved in T2D^3, 9^.

In this work we construct a regulatory network (Figure 1) based on a bottom-up approach that integrates existing knowledge of essential relations of pancreatic-beta cell components leading to MS and T2D (an alternative methodology contemplated in other works is a top-down approach based on inference algorithms to reconstruct the network topology^10^). A network is a system formed by connected elements, called nodes, and relations between them. Every node represents a gene, a transcription factor, a cytokine, etc. The network dynamics can be studied by mathematical models to analyze steady states, also called attractors, which define expression patterns associated to a cellular fate. The use of these models may increase the understanding of the cellular differentiation process or even a disease evolution^11–14^. In this context, Waddington introduced in 1957 the concept of epigenetic landscape to study the evolution of the cells as a phenomenon driven by the landscape dynamics. He proposed a metaphor of the epigenetic landscape where the cellular development is viewed as a ball rolling down in a landscape formed by peaks and valleys^15^. Following its trajectory, the ball may finally fall into a valley, representing its final position. This position can be associated with the concept of attractor and represents a cellular equilibrium condition. The network dynamics can be analyzed studying the related landscape^16–18^.

**Figure 1.**
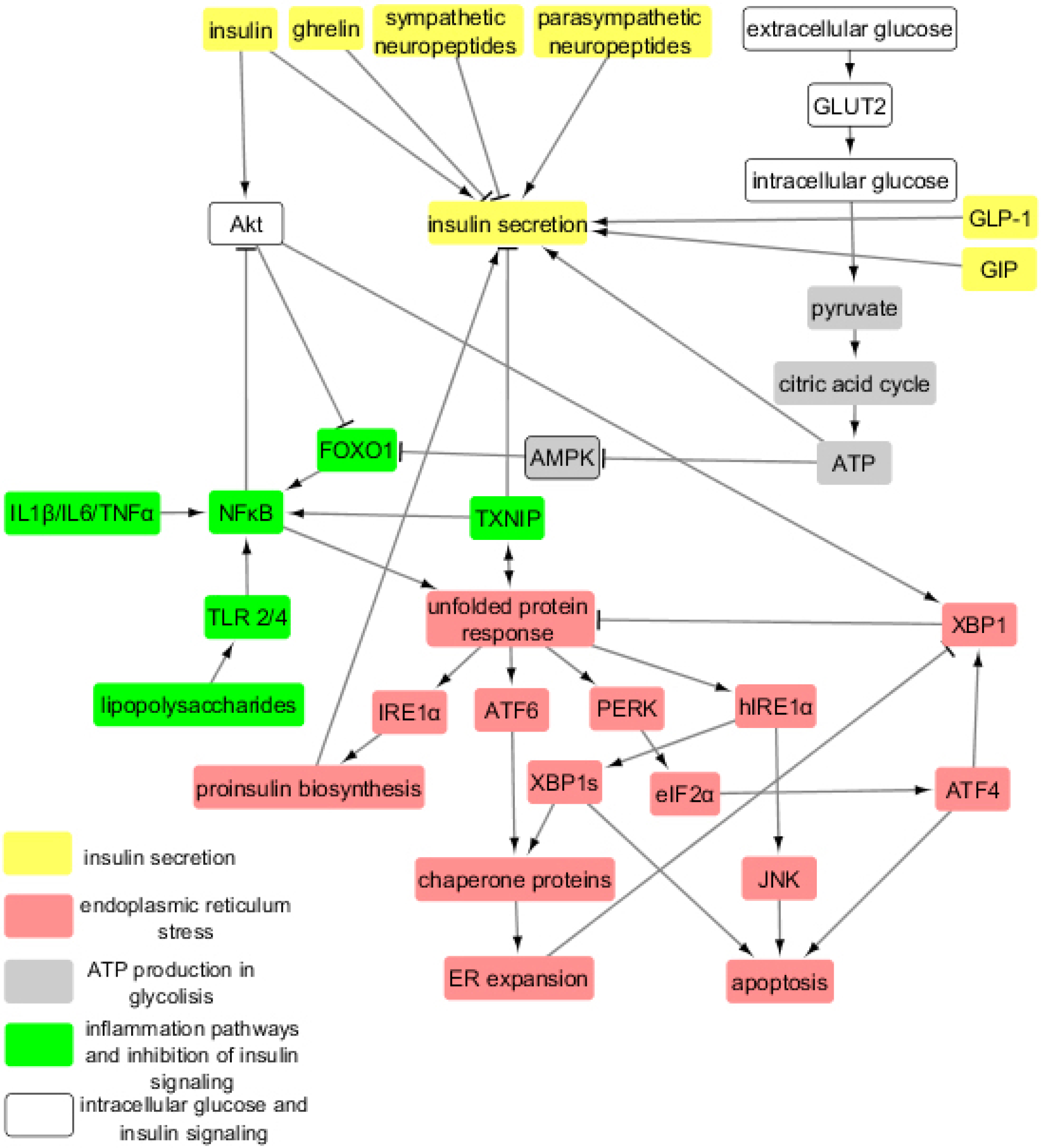
Interaction network in pancreatic-beta cells. The network include cytokines, proteins, hormones, receptors, transcription factors, etc., which are central in the evolution of health to metabolic syndrome, and type 2 diabetes. The colors of the rectangles represent processes and signaling pathways altered during the T2D evolution. The arrows and bars correspond to positive and negative regulatory interactions, respectively.

Some diseases like cancer have been studied as a process with an attractor associated to health and its transition to another cell fate corresponding to a cancer attractor associated to aberrant gene expression profiles^19–21^. In this regard, the balance between robustness and plasticity in biological systems plays a prime role. Robustness describes the insensitivity of a system to external perturbations, such as accumulated mutations in cancer; plasticity describes the changeability of a system in response to modifications in micro-environment conditions. In the present work we describe the progression to T2D based on the plasticity mechanism, in other words, in the impairment of metabolic processes due to alterations of exogenous and endogenous environment of pancreatic-beta cells. With that purpose, the network includes stimulatory and inhibitory interactions responsible of glucose internalization, processing, and storage. The dynamic evolution of these relations generates processes that with the course of time modify homeostatic expression levels of the network components, eventually leading to an alternative equilibrium regime determined by the particular set of the system initial conditions. Any equilibrium state or attractor corresponds to a pattern of activation or inhibition of the cellular components, which may display either a fixed-point or cyclic character. We identify these expression patterns with states of health, MS, or T2D.

A particular approach to model the network interactions is the Boolean network, fist proposed by Stuart Kauffman in 1960s as a mathematical model of genetic regulation^22^. In this scheme, it is assumed that the genes can only take two values related to activated (1), or inhibited states (0); these values depend on previous states of all the genes in the network. Thus, by considering a network formed by *n* genes, the state of the gene *k* at a time *t* + *τ* can be described by the mapping *q*_*k*_(*t* + *τ*) = *f*_*k*_(*q*_1_(*t*), …, *q*_*n*_(*t*)). Here, *f*_*k*_ is determined by a logical proposition that satisfies Boole axiomatics. Kauffman′s hypothesis establishes that the stationary states of the mapping determined by the condition *q*_*k*_(*t* +*τ*) = *q*_*k*_(*t*) define different cellular phenotypes. The network dynamics based on Boolean models can give us an idea of all the stationary states in a biological system. This approach has been formerly employed to characterize the underlying mechanistic for a number of biological processes^18, 21, 23–29^.

Boolean models provide important information on the basic relations architecture that determine alternative cell fates, and can be used for the analysis of biological circuits without requiring explicit values of the network parameters^10^. In this context, we study a regulatory network of 35 pancreatic-beta cell components supported on a review of current experimental and clinical evidence (Table 1). The network comprises connected sub-networks representing insulin signaling and glucose internalization, adenosine triphosphate (ATP) production in glycolisis, inflammation pathways, endoplasmic reticulum stress, and insulin secretion (Figure 1). The network dynamics provided a number of steady states that were broadly classified into three main groups consistent with health, MS, and T2D (Table 2). The progression of the disease was described as a transit between these later stages. With that purpose, we introduced a more realistic approach that considers that the expression levels, concentrations and parameters of the system may acquire any real value with a continuous range^18^. The approach is based on fuzzy logic, originally aimed to provide formal foundation to approximate reasoning^30–33^. In this scheme, the truth values of variables may vary from completely false (0) to completely true (1), and the degree to which an object exhibits a property *p* is given by a characteristic function *μ*[*p*] with continuous variation in the range {0, 1}.

**Table 1.** Experimental evidence of each node and its function in the network.

| Node | Experimental evidence |
| --- | --- |
| **Insulin secretion** |  |
| Insulin | Leibiger |
| Parasympathetic neuropeptides | Chiu,Ahren2,Koh |
| GLP-1 | Velasco,Phillips |
|  | Baggio,Ahren |
| GIP | Velasco,Phillips |
|  | Baggio,Ahren |
| Ghrelin | Tong,Pradhan |
| Sympathetic neuropeptides | Chiu,Ahren2,Koh |
| Insulin secretion | Velasco,Phillips |
|  | Baggio,Ahren |
|  | Chiu,Ahren2 |
|  | Koh,Tong |
|  | Pradhan,Leibiger |
| **Intracellular glucose and insulin signalling** |  |
| Extracellular glucose | Prentki2,Thorens,Stolarczyk |
| GLUT2 | Prentki2,Thorens,Stolarczyk |
| Intracellular glucose | Prentki2,Thorens,Stolarczyk |
| Akt | Tuttle,Kops,Cras-Meneur |
| **Glycolysis and ATP production** |  |
| Pyruvate | Muoio,Hiriart |
|  | Velasco2,Nolan |
| Citric acid cycle | Muoio,Hiriart |
|  | Velasco2,Nolan |
| ATP | Muoio,Hiriart |
|  | Velasco2,Nolan |
| AMPK | Muoio,Hiriart |
|  | Velasco2,Nolan |
| **Inflammation pathways** |  |
| IL1β/IL6/TNFα | Donath,Hotamisligil,Vives-Pi |
| TLR 2/4 | Garay-Malpartida,Vives-Pi |
|  | Guilherme,Laat,Jayashree |
| Lipopolysaccharides | Garay-Malpartida,Vives-Pi |
|  | Guilherme,Laat,Jayashree |
| FOXO1 | Fang,Kitamura,Kops |
| NFκB | Shoelson,Hotamisligil |
| TXNIP | Andrews,Lerner |
| **Endoplasmic reticulum stress** |  |
| Unfolded protein response | Chen,Rabhi,Hetz |
| XBP1 | Chen,Rabhi,Hetz |
| IRE1α | Chen,Rabhi,Hetz |
| Proinsulin biosynthesis | Chen,Rabhi,Hetz |
| PERK | Chen,Rabhi,Hetz |
| eIF2α | Chen,Rabhi,Hetz |
| ATF4 | Chen,Rabhi,Hetz |
| ATF6 | Chen,Rabhi,Hetz |
| hIRE1α | Chen,Rabhi,Hetz |
| XBP1s | Chen,Rabhi,Hetz |
| Chaperone proteins | Chen,Rabhi,Hetz |
| ER expansion | Chen,Rabhi,Hetz |
| JNK | Chen,Rabhi,Hetz |
| Apoptosis | Chen,Rabhi,Hetz |

**Table 2.** Expression patterns associated to health, metabolic syndrome, and T2D. The state of inhibition or activation of each of beta cells nodes are represented by 0 o 1, respectively.

| Node | Definition | Health state | MS state | T2D state |
| --- | --- | --- | --- | --- |
| **Insulin secretion** |  |  |  |  |
| Insulin | Insulin | 1 | 1 | 0 |
| Parasympathetic neuropeptides | Parasympathetic neuropeptides | 1 | 1 | 0 |
| GLP-1 | Glucagon-like peptide 1 | 1 | 1 | 0 |
| GIP | Glucose-dependent<br>insulinotropic peptide | 1 | 1 | 0 |
| Ghrelin | Ghrelin | 0 | 0 | 1 |
| Sympathetic neuropeptides | Sympathetic neuropeptides | 0 | 0 | 1 |
| Insulin secretion | Insulin secretion | 1 | 1 | 0 |
| **Intracellular glucose**<br>**and insulin signalling** |  |  |  |  |
| Extracellular glucose | Extracellular glucose | 1 | 1 | 1 |
| GLUT2 | Glucose transporter 2 | 1 | 1 | 1 |
| Intracellular glucose | Intracellular glucose | 1 | 1 | 1 |
| Akt | Protein kinase B | 1 | 1 | 0 |
| **Glycolysis and ATP**<br>**production** |  |  |  |  |
| Pyruvate | Pyruvate | 1 | 1 | 1 |
| Citric acid cycle | Citric acid cycle | 1 | 1 | 1 |
| ATP | Adenosine triphosphate | 1 | 1 | 1 |
| AMPK | AMP-activated protein kinase | 1 | 0 | 0 |
| **Inflammation pathways** |  |  |  |  |
| IL1β/IL6/TNFα | Interleukin 1 beta/<br>Interleukin 6/Tumor necrosis factor | 0 | 1 | 1 |
| TLR 2/4 | Toll-like receptor 2 or 4 | 0 | 1 | 1 |
| Lipopolysaccharides | Lipopolysaccharides | 0 | 1 | 1 |
| FOXO1 | Forkhead box protein O1 | 0 | 1 | 1 |
| NFκB | Nuclear factor kappa-light<br>-chain-enhancer of activated B cells | 0 | 1 | 1 |
| TXNIP | Thioredoxin interacting protein | 0 | 1 | 1 |
| **Endoplasmic reticulum**<br>**stress** |  |  |  |  |
| Unfolded protein response | Unfolded protein response | 0 | 1 | 1 |
| XBP1 | X-box binding protein 1 | 1 | 1 | 0 |
| IRE1α | Inositol-requiring enzyme 1 | 1 | 1 | 0 |
| Proinsulin biosynthesis | Proinsulin biosynthesis | 1 | 1 | 0 |
| PERK | Protein kinase R-like<br>endoplasmic reticulum kinase | 0 | 0 | 1 |
| eIF2α | Eukaryotic initiation factor 2 | 0 | 0 | 1 |
| ATF4 | Activating transcription factor 4 | 1 | 1 | 1 |
| ATF6 | Activating transcription factor 6 | 0 | 0 | 1 |
| hIRE1α | Hyperphosphorylation of IRE1α | 0 | 0 | 1 |
| XBP1s | X-box binding protein 1 splicing | 0 | 0 | 1 |
| Chaperone proteins | Chaperone proteins | 0 | 0 | 1 |
| ER expansion | Endoplasmic reticulum expansion | 0 | 0 | 1 |
| JNK | C-Jun N-terminal kinase | 0 | 0 | 1 |
| Apoptosis | Apoptosis | 0 | 0 | 1 |

The transit between different disease stages was characterized by merging the fuzzy network approach with the general theory of non-equilibrium phase transitions and self-organization^34, 35^. According to this theory, cooperative phenomena arise from non-linear interactions of a large number of elementary subsytems, leading to the emergence of ordered patterns or phases. The theory relies upon two main concepts: the notion of order parameter and the adiabatic elimination of irrelevant system variables based upon a hierarchy of time constants present in most systems. The order parameters are those variables that mainly drive the system dynamics. Their relaxation times are usually much longer than those of irrelevant variables and work like control parameters of the system. Irrelevant variables decay rapidly to a steady state, so that they may be effectively eliminated from the overall dynamics. In this view, the order parameters define the general features of the system, including the final expression patterns associated to a set of initial conditions, while less relevant variables adapt their values to the instructions dictated by the order parameters. It may be shown that transitions between different ordered phases may be induced by modifications of the control parameters. In Waddington’s landscape context, this mechanism may be interpreted as alterations of the peaks separating the valleys, allowing the exploration of the landscape and transit between valleys.

In our study, we assumed that the phases corresponding to health, MS and T2D were those already determined by the Boolean approach and studied the possible phase transitions between them by considering a set of ordinary differential equations for the expression levels of the fuzzy network components^18, 27, 36, 37^: 
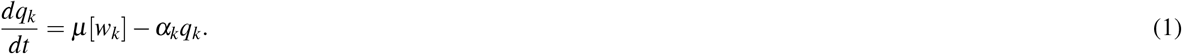

## 1 Materials and methods

### 1.1 Network construction

A network of 35 elements for pancreatic-beta cells regulation was constructed based on a literature review (Table 1). With that purpose, we first identified connected subnetworks that constitute key elements participating in glucose intake, processing, and storage. These processes are associated with insulin signaling and secretion, glucose internalization, glycolysis and adenosine triphosphate (ATP) production, inflammatory pathways, and endoplasmic reticulum (ER) stress. The key elements conforming each subnetwork and their interactions are briefly described below.

#### 1.1.1 Intracellular glucose and insulin signaling

In pancreatic-beta cells, the glucose-stimulated insulin secretion (GSIS) is a consequence of increased circulating glucose concentrations, and the glucose transporter 2 (GLUT2) participates in this process. GLUT2 leads to an equilibrium between extra and intracellular glucose^38, 39^. In different animal models of diabetes, it has been proposed that if the expression of GLUT 2 is strongly reduce, impaired GSIS is associated whit this fact, whilst GLUT 2 is active in the control of GSIS^40^.

In addition to extracellular glucose, peripheral insulin may stimulate insulin secretion in pancreatic-beta cells. The insulin receptor signaling includes the activation of phosphoinositol 3-kinase (PI3K) and the subsequent expression of protein kinase B (PKB), also known as Akt. The over-expression of PKB/Akt has been related to mass expansion and insulin secretion increase of Langerhans islets, while its under-expression results in reduced islet mass and impaired pancreatic-beta cell function^41^. An overexpression of active Akt in mouse pancreatic-beta cells models has been associated to improving glucose tolerance and resistance to experimental diabetes, being Akt central in pancreatic-beta cells size and function^42, 43^.

#### 1.1.2 Glycolysis and ATP production

When extracellular glucose increases, it is internalized in pancreatic-beta cells through a GLUT2 transporter. Glucose then enters into the glycolysis cycle: The associated metabolic processes includes mitochondrial oxidation, which generates and increases the ratio ATP/adenosine diphosphate (ADP). This change in ATP/ADP relationship closes the ATP sensitive potassium (K) channels, and the membrane depolarizes activating the influx of calcium (Ca), resulting in insulin exocytosis. A model of MS induced in adult male Wistar rats with high sucrose diet after 24 weeks of treatment showed central obesity, hyperinsulinemia, insulin resistance and an increased sensitivity to ATP in ATP sensitive K channels, suggesting that this change can explain the increase in insulin secretion^44–47^. This is the course followed by the major triggering pathway for GSIS^48^. Insulin interacts with their targets by binding to the insulin receptor and producing tyrosine transphosphorylation of its internal subunits, and a cascade of protein phosphorylation, starting with the substrate of the insulin receptor proteins (IRS-1 and 2).

#### 1.1.3 Insulin secretion

Several factors regulate insulin secretion by pancreatic-beta cells; the primary are nutrients, mainly glucose. Others are some amino acids, free fatty acids, hormones and neurotransmitters. For example, some insulin potentiators are the incretins, glucagon-like peptide 1 (GLP-1) and glucose-dependent insulinotropic peptide (GIP). They are produced in the intestine and increase the GSIS. In vitro and in preclinical models, GLP-1 and GIP promote pancreatic-cell proliferation and potentiate nutrient-induced insulin secretion^5, 49, 50^. It has been corroborated that the insulin secretory response to GLP-1 is augmented in insulin-resistant mice compared with normal mice^51^.

The hormone ghrelin is produced during fasting by the stomach and inhibits the GSIS, probably by opening the ATP-sensitive K channels and inhibiting the activation of Ca channels. Studies of ghrelin infused in healthy humans have shown to worsen the glucose and pancreatic-beta cell responses to meal ingestion, ghrelin decreased insulin sensitivity, impaired pancreatic-beta cell function, and induced glucose intolerance^52, 53^.

Neuropeptides are released molecules involved in neural communication; some of them also regulate insulin secretion and protect the brain from possible damages due to hypoglycemia; parasympathetic neuropeptides, released during electrical vagal activation like vasoactive intestinal polypeptide (VIP) stimulate insulin secretion, while sympathetic islet neuropeptides inhibit insulin secretion^54^. Several rodent models of diabetes have proved the stimulatory effect of parasympathetic nerves and the inhibitory action of sympathetic nerves over insulin secretion^55, 56^.

It has been reported by mice models that secreted insulin also stimulates the immediate process of insulin exocytosis, the probable explanation is due to an insulin-dependent increase in cytosolic free Ca^57^.

#### 1.1.4 Inflammation pathways

High levels of glucose and free fatty acids can stimulate the release of pro-inflammatory cytokines, for example, interleukin 1*β* (IL1*β*), interleukin 6 (IL6), and tumor necrosis factor (TNF). These molecules produce inflammation of pancreatic-beta cells, impairment of insulin secretion, and apoptosis^8^. In obese mouse models, a lack of TNF function has resulted in improved insulin sensitivity and glucose homeostasis. Cytokines may activate kinases that participate in serine phosphorylation of the insulin receptor or the IRSs, modifying insulin action, activating, for example, the nuclear factor kappa-light-chain-enhancer of activated B cells (NF*κ*B)^58^. The antiinflammatory actions of the thiazolidinediones (TZDs) have shown inhibit NF*κ*B and improve insulin signaling^59^.

Toll-like receptors 2 and 4 (TLR 2/4) are cell-surface receptors that activate the innate immune response and recognize a wide variety of antigenic motifs like lipopolysaccharides (LPS) and lipoproteins of bacterial walls. However, they may be also triggered by excess of circulating fatty acids, inducing modifications in insulin signaling, persuading inflammation^60–62^. Pancreatic-beta cell line MIN6 have shown increase of TLR 4 in response to LPS and a decreased insulin synthesis and secretion^63, 64^.

Thioredoxin interacting protein (TXNIP) is a protein that play an important role in the type 2 diabetes (T2D) development, activates reactive oxygen species and cytokines and its over expression induces ER stress and pancreatic-beta cell apoptosis^65, 66^.

One of the forkhead transcription factor class O isoforms, (FoxO1), is the most abundant isoform in pancreatic-beta cells and regulates its immediate target gene p27 at the mRNA level. It has been exhibited in cell lines of rat insulinoma INS-1 and in mammalian cell lines that PI-3K activates Akt through phosphorylation, and active Akt, in turn, inhibits the transcriptional activation of FoxO1 via phosphorylation-dependent nuclear exclusion, resulting in inhibition of its target gene expression. On the other hand, excessive FOXO1 activation has shown irreversible apoptosis in pancreatic-beta cells^43, 67, 68^.

#### 1.1.5 Endoplasmic reticulum stress

When the folding capacity of the ER is exceeded, misfolded or unfolded proteins accumulate in the ER lumen, resulting in ER stress. During the T2D progression, pancreatic-beta cells produce more insulin in response to elevated levels of glycaemia and insulin resistance, where the ER regulates its capacity in order to prevent unfolded proinsulin accumulation and to preserve the ER homeostasis. These signaling pathways are designated as the unfolded protein response (UPR). There have been genetic and biochemical studies on mouse models and human UPR sensor mutations that describe the processes involved in the UPR response during the progression of T2D to prevent pancreatic-beta cell failure and the disease adaptation^69^.

Under normal physiological conditions UPR reduces ER stress triggering transcription of folding proteins, attenuates eukaryotic initiation factor 2 (eIF2*α*) protein translation, and promotes proinsulin synthesis via inositol-requiring enzyme 1 (IRE1*α*) kinase. On the other hand, under physiopathological conditions IRE1*α* may activate several downstream signaling of the UPR. For example, by splicing the X-box binding protein 1 (XBP1) mRNA, to enhance the protein-folding capacity, or triggering apoptosis when the ER recovery mechanism fails. Similar to IRE1, the protein kinase R-like endoplasmic reticulum kinase (PERK) and the activating transcription factor 6 (ATF6) are ER transmembrane proteins that contain an ER luminal stress-sensing domain. PERK may inhibit the increase in protein-folding demand by phosphorylating eIF2*α*, which can also activate the activating transcription factor 4 (ATF4) to regulate UPR target genes. In mammals, PERK-eIF2*α*-ATF4 activate ER stress-triggered apoptosis^70, 71^. High levels of ER stress also activate the C-Jun N-terminal kinase (JNK) by the kinase activity of IRE1 and participate in the apoptosis process^69^. ATF6 and XBP1s drive to expression of ER chaperon and ER expansion. Experiments with ATF6 knockout mice fed with a high-fat diet to create diet-induced obesity have demonstrated a severe hypoglycaemia suggesting that suppression of ATF6 increased insulin sensitivity^72^. In the present work, we will assume that besides its normal action, UPR concurrently displays elements associated to physiopatological conditions.

### 1.2 Steady states: Boolean analysis

The stationary states of a Boolean network with *n* nodes are determined by the final steady states resulting from the dynamical trajectories of the system with origin at a set of 2^*n*^ possible initial conditions, associated to a state of total expression (*q*_*i*_(*t* = 0) = 1), or total inhibition (*q*_*i*_(*t* = 0) = 0) of every component of the network. However, for a network with *n* = 35 nodes considered in this study, this is an exceedingly large number (more than 34 billions) of different configurations. In order to circumvent this difficulty, we divided the node interactions into two sets, corresponding to independent and dependent interactions. In the first set we considered those that are external inputs, as well as those that due to their centrality, determine the states of other nodes through the Boolean interactive rules. In the input set we identified nine components: extracellular insulin, ghrelin, sympathetic neuropeptides, parasympathetic neuropeptides, extracellular glucose, IL1*β*/IL6/TNF*α* (inflammatory cytokines), lipopolysccharides, GLP-1, and GIP. To this set we also added Akt and UPR due to their strategic positions within the network. They both connect circuits involved in the endoplasmic reticulum stress and the inflammatory response, so that their expression levels influence the pancreatic beta-cell fates through alternative signaling routes. Although these nodes involve dependent interactions, they were considered as inputs subject to the condition that the rest of variables actualize their values consistently with the Boolean rules. In particular, we contemplated that since Akt is controlled upstream by insulin, if this latter element is inactive then Akt is also inactive; on the other hand, if insulin is active then Akt may initiate with any value 0 or 1. It follows that the analysis can be reduced to consider only a set of 2^11^ = 2048 initial conditions.

The Boolean analysis was carried up by means of a program in Fortran 90 (Supplementary material 1) that provides the attractors of the dynamical system (Supplementary material 2). Briefly, the code creates two arrays, the first one corresponds to a matrix of all the possible configurations of initial conditions and the second one is formed by a matrix with all the possible configurations of final conditions, one for each initial configuration. The code specifies the number *N*_*k*_ and the specific initial conditions that lead to the same attractor *k*. *N*_*k*_ determines the size of the attraction basin and provides, in principle, an estimation of the probability that a given activation and inhibition pattern is expressed. Since the 2^11^ configurations of initial conditions are generated independently, each initial configuration would entail an *a priori* probability *p* = 1/2^11^ that a given attractor *k* is reached, so that the probability that the *k*-th pattern is expressed would be *P*_*k*_ = *N*_*k*_*p*. However, certain combinations of input variables could have no realistic biological sense, and such combinations should be discarded in a probabilistic counting.

In order to classify the steady states of the network into states corresponding to health, MS, and T2D stages, we constructed Table 2, based on a review of current literature. This table represents a kind of ‘idealistic’ classification scheme since, due to the multifactorial nature of the disease, any specific patient may show a hybrid pattern of the elements considered in the table, which may besides evolve at different rates with the progression to TD2.

From the mathematical point of view, it is also unlikely to achieve exact coincidence with the ideal values corresponding to health, MS, and T2D in Table 2, since the regulatory network includes a relatively large number of nodes with interaction rules that are not heavily restrictive as compared, for example, with Boolean rules describing cell differentiation pathways^18^. Therefore, in Supp. Mat. 2 we considered classification criteria based on the following: There are four fundamental nodes that determine the former states: insulin secretion, XBP1, TXNIP, and apoptosis. For health, it is necessary that the nodes corresponding to insulin secretion, and XBP1 are active, while those corresponding to TXNIP and apoptosis are inactive; in addition, the modules corresponding to glucose internalization and ATP production are active, while the modules corresponding to inflammation and endoplasmic reticulum stress are mostly inactive. For MS, the node corresponding to insulin secretion is active, while that corresponding to apoptosis is inactive: on the other hand, XBP1 and TXNIP may be either active or inactive; as before, the modules corresponding to glucose internalization and ATP production are active, but now the module corresponding to inflammation is mostly active. For TD2, it is necessary that the nodes corresponding to insulin secretion, and XBP1 are inactive, while those corresponding to TXNIP and apoptosis are active; in addition, the modules corresponding to inflammation and endoplasmic reticulum stress are mostly active.

### 1.3 Continuous logic approach

Fuzzy logic propositions comply with Boolean axiomatics (except for the excluded-middle law), and they may be obtained from the original Boolean propositions *f* by replacing Boolean operators by fuzzy connectors (see Supplementary Material 3 and 4). The translation *f* (*q*_1_, …, *q*_*n*_) → *w*(*q*_1_, …, *q*_*n*_) may be performed by considering the following rules: *q* and *p* → *q·p*, *q* or *p → q* + *p−q·p*, not *p* → 1−*p*. Thus, for example, the Boolean proposition *f*_*k*_ = *q*_1_ or *q*_2_ and not *q*_3_, becomes *w*_*k*_ = (*q*_1_ + *q*_2_−*q*_1_·*q*_2_)(1−*q*_3_). In this work, the characteristic function is expressed by means of a function with a sigmoid structure: 

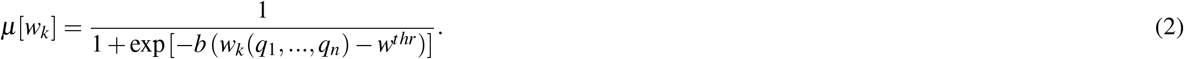

Here, *w*^*thr*^ is a threshold value that renders *w*_*k*_ true (expressed) if *w*_*k*_ > *w*^*thr*^. The simplest assumption is to consider *w*^*thr*^ = 1/2. The parameter *b* is a variation rate for the change of the proposition *μ*(*w*_*k*_) from an unexpressed to a totally expressed state, being gradual for small *b*, and steep for large *b*. In the limit *b* ≫ 1, the change is sudden, similarly as in the Boolean description. It can be shown that the results of the model are independent of the choice *b* as far as this parameter is large enough. In this work we suppose that *b* = 5.

### 1.4 Transitions between steady states

Dynamical analyses were performed to study the transitions from health to a MS stage, and from a MS stage to T2D, by employing the continuous regime. this was analyzed with Wolfram Mathematica computing system, using ordinary equations described in the mathematical model section afore mentioned see Supplementary Material 5). Each attractor in Table 2 was in this step considered as a set of initial conditions of every node, and the analysis allowed to identify cases where the perturbation of a node decay rate induced a transition from the original to a different attractor, it was initially assumed that *α*_*k*_ = 1 for every node. To address this goal, every decay rate was gradually increased from its initial value *α*_*k*_ = 1 to a maximal value *α*_*k*_ = 5, allowing to determine its capacity to generate a qualitative modification of the system dynamics. This is equivalent to considering exhaustion mechanisms of cellular components, the action of overnutrition, chronic inflammation, continuous ER stress, etc.

In this regard, for the simulation of health to MS, we took as initial conditions the health stage shown in table 1 and changed the decay rate of every active node to find out the main factors responsible from the shift between health and MS. The same process occurred to identify the principal factors involved in the transition between health to a transient MS state, and a final T2D stage. We also tested a reversible behavior from an advanced disease stage to an improved one.

## 2 Results

### 2.1 Steady states related to health, metabolic syndrome, and type 2 diabetes

As described in the Methodology section (Boolean analysis), even though the network consists of 35 nodes involving 2^35^ possible initial conditions, the totality of steady states were determined by considering only 2^9^ initial conditions given by 9 external input nodes, times 2^2^ initial conditions associated to 2 central internal nodes, leading to a total number of 2^11^ initial conditions. From each of these conditions the system evolved into final states defining specific activation and inhibition patterns. They were classified according to their congruence with characteristic patterns associated to health, metabolic syndrome (MS), or type 2 diabetes (T2D), depicted in Table 2. For example, stages defined as health and or MS showed a variable expression of inflammatory factors, but inactive apoptosis, while the T2D group displayed apoptosis and expression of thioredoxin interacting protein (TXNIP), but inhibition of insulin secretion and X-box binding protein 1 (XBP1). From the total set of 2048 stationary states, 32 corresponded to health, 186 to MS, and 1024 to TD2 stages.

An analysis of the network architecture in Figure 1 performed by collapsing downstream nodes along a given pathway let us identify fundamental functional modules involved in the disease. In Figure 2 we observe that the transcription factor TXNIP, a suppressor of insulin secretion, plays a central role in the description of T2D, acting like a bridge between the endoplasmic reticulum (ER) stress and the inflammatory response. It also constitutes a feedback circuit with the nuclear factor kappa-light-chain-enhancer (NF*κ*B), and the unfolded protein response (UPR), which may conduce to a vicious deleterious circle under physiopathological conditions. In turn, the UPR defines a switching module with protein kinase B (AkT) through the action of the transcription factor XBP1. In the case where UPR is activated, it can activate XBP1 by the alternate way showed in dotted lines and also has the capacity to suppress it by the direct way. A switching module is defined by a pair of mutually inhibitory interactions, so that the preponderance of one or the other drives alternative signaling pathways.

**Figure 2.**
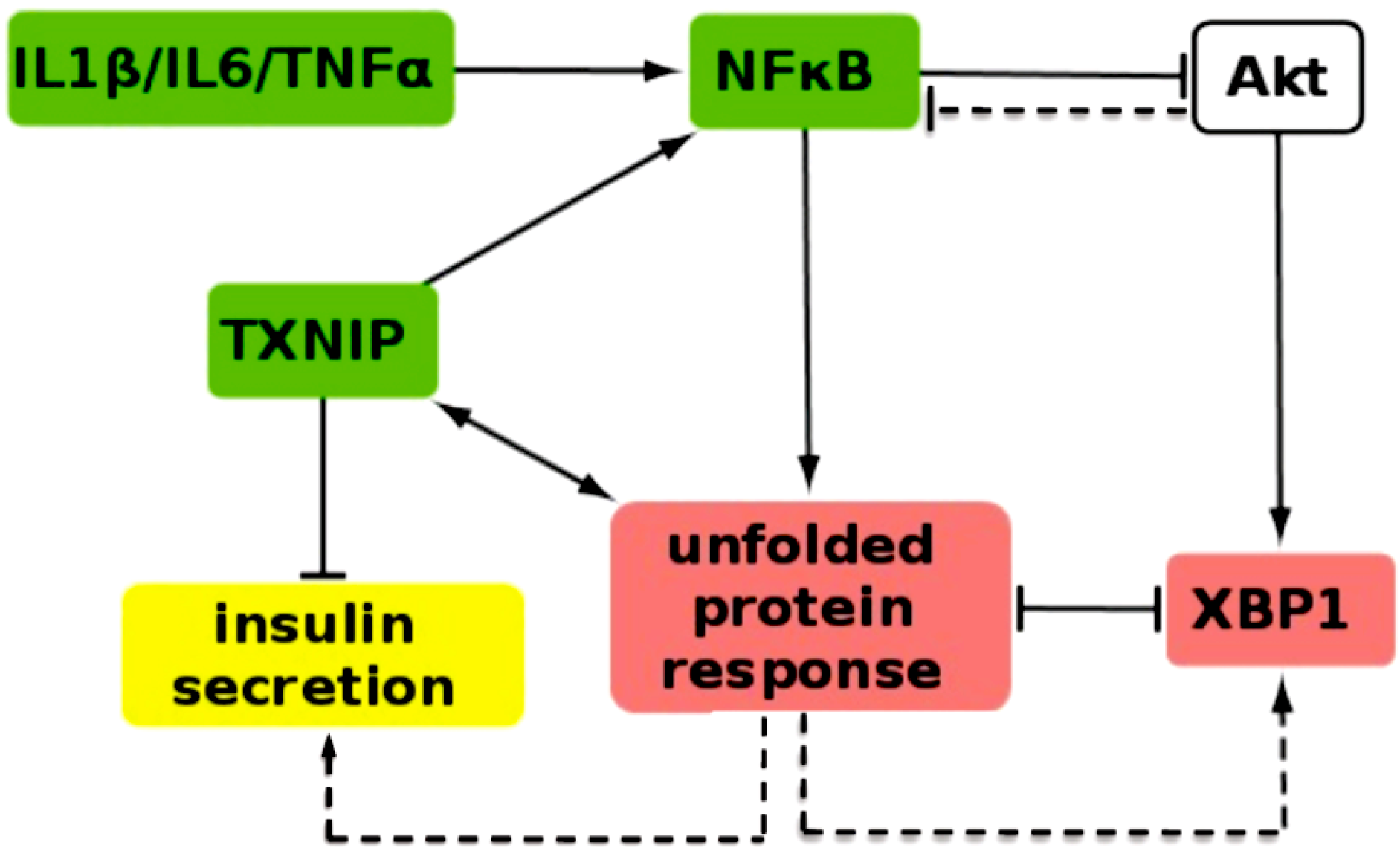
Central module connecting insulin signaling, ER stress, inflammation, and insulin secretion suppression. TXNIP acts as a central bridge between the three later responses. XBP1 and UPR form a switching module resulting from mutual inhibition.

### 2.2 From health to metabolic syndrome

In Figure 3 we observe an overall dynamic behavior consistent with a transition from health to MS, originated by an incremented decay rate of the signaling molecule Akt. This alteration promotes inflammatory factors like the nuclear factor NF*κ*B, and the forkhead box protein *O*1 transcription factor transcription factor (FOXO1). In addition, insulin secretion decreases from normal levels displaying an oscillatory behavior, while the UPR is stimulated, together with the transcription factor TXNIP that acts like a bridge between inflammatory and endoplasmic reticulum (ER) stress pathways. The former predictions are consistent with experimental evidence describing Akt as a factor that regulates the expression of gluconeogenic and lipogenic enzymes by controlling the activity of the FOXO1, which in the network promotes the action of NF*κ*B.

**Figure 3.**
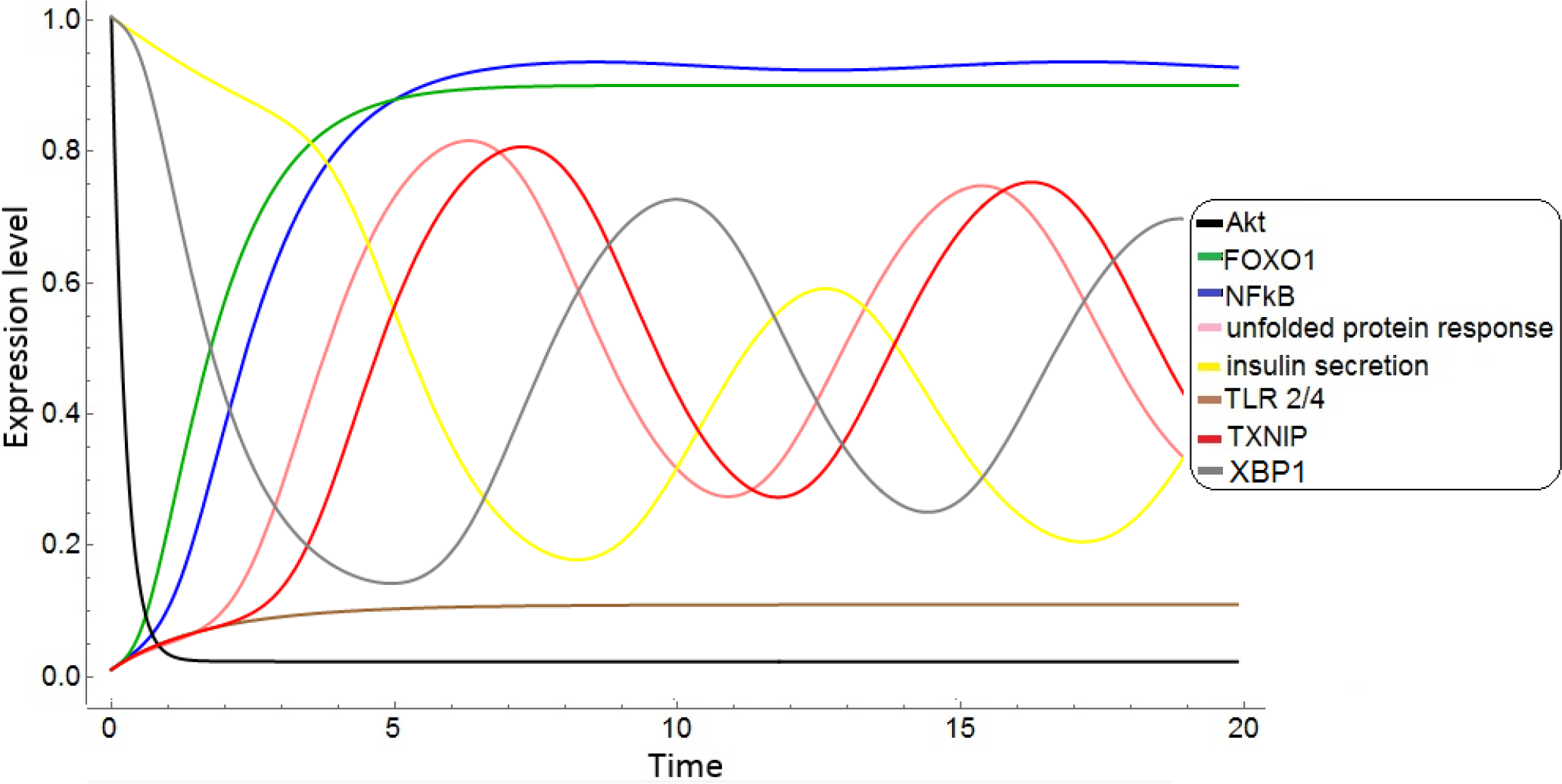
From health to metabolic syndrome. Dynamic evolution of pancreatic-beta cells expression profiles associated to under-expression or exhaustion of two key cellular components involved in metabolic regulation: AkT and insulin. The healthy state is associated to normal expression of the nodes AkT, insulin secretion, and XBP1, with a null initial expression of the rest of nodes depicted in the figure. The decay rates of all nodes in the beta cells network are assumed *α*_*k*_ = 1, except for insulin and Akt, suppressed at increased rates *α*_*ins*_ = *α*_*Akt*_ = 2, respectively. As a result, a strong increment of NF*κ*B, and FOX01 arise. UPR and TXNIP show oscillations about an increased mean level, while XBP1 and insulin secretion show a delayed oscillatory behavior around a reduced mean value.

We observe that as a consequence of the inhibition of Akt, several elements display oscillations of their expression levels. These can be explained as a consequence of the design of the network. Some highly connected nodes are subject to the simultaneous action of positive and negative regulatory interactions which create effective feedback loops. As is apparent in Figure 2, if Akt decays, it suppresses XBP1 and then UPR is activated, inducing the expression of TXNIP and NFkB, thus creating a feedback loop UPR → TXNIP → NFkB → UPR which suppresses insulin secretion. In turn, UPR also activates XBP1 by an alternatative pathway (in dotted lines) leading to inhibition of the UPR loop, thus promoting insulin secretion but also inhibiting XBP1 again, so that an oscillatory cycle ensues.

### 2.3 From health to transient metabolic syndrome, and final manifest diabetes

In Figure 4 we display an alternative pathway from health to a transient MS state, and a final T2D stage generated by a increased decay rate of the transcription factor XBP1. The initial condition is a healthy state with normal Akt and insulin secretion levels, but no ER stress or inflammatory symptoms. The increase of the decay rate of XBP1, drives the transition from health to an intermediate MS stage, and a final transition to T2D. In this situation, NF*κ*B, FOX01, TXNIP, and UPR show a gradual expression increase, while Akt and insulin secretion display a steady decrease along time. The intermediate states between health and T2D may be interpreted as associated to an evolving MS stage with progressive symptomatology.

**Figure 4.**
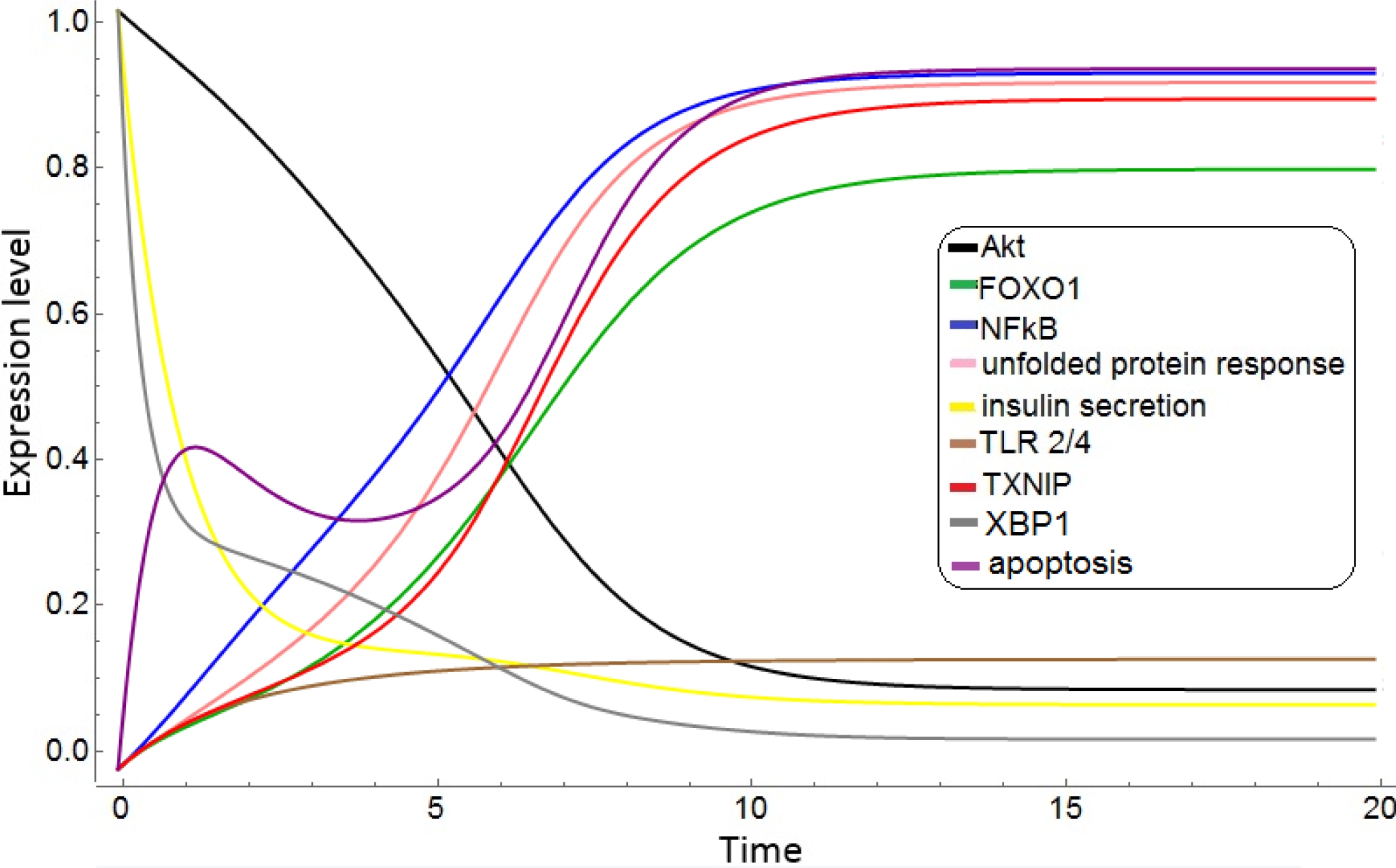
From health to transient metabolic syndrome, and subsequent type 2 diabetes. Dynamic evolution of pancreatic-beta cells expression profiles associated to under-expression of XBP1. The healthy state is associated to normal expression of AkT, insulin secretion, and XBP1, with a null initial expression of the rest of nodes depicted here. The decay rates of all nodes in the beta cells network are *α*_*k*_ = 1, except for the transcription factor XBP1, with an increased decay rate of *α*_*XBP1*_ = 3. With the course of time, the system transits from the healthy state, to a transient metabolic syndrome state where all considered factors display intermediate levels. Finally, the system reaches a final stage of manifest diabetes, where Akt, insulin secretion, and XBP1 are strongly suppressed, while endoplasmic reticulum stress, inflammatory signals, and apoptosis are over-expressed.

### 2.4 Improvement of symptomatology

Figure 5 describes a reverse transition from an advanced disease stage to an improved stage. This is induced by an increased decay rates of TXNIP, as well as of exogenous LPS. In that case, insulin secretion is recovered from an initial impaired condition to a state with quasi-normal levels. On the other hand, inflammatory signals still persist showing a similar behavior as that depicted in Figure 3. Notably, Akt acquires a low-level but finite expression, in contrast with the case represented in Figure 3 where it is completely inhibited. This suggests that therapies targeting TXNIP expression accompanied by caloric restrictions could improve symptomatic manifestations of T2D.

**Figure 5.**
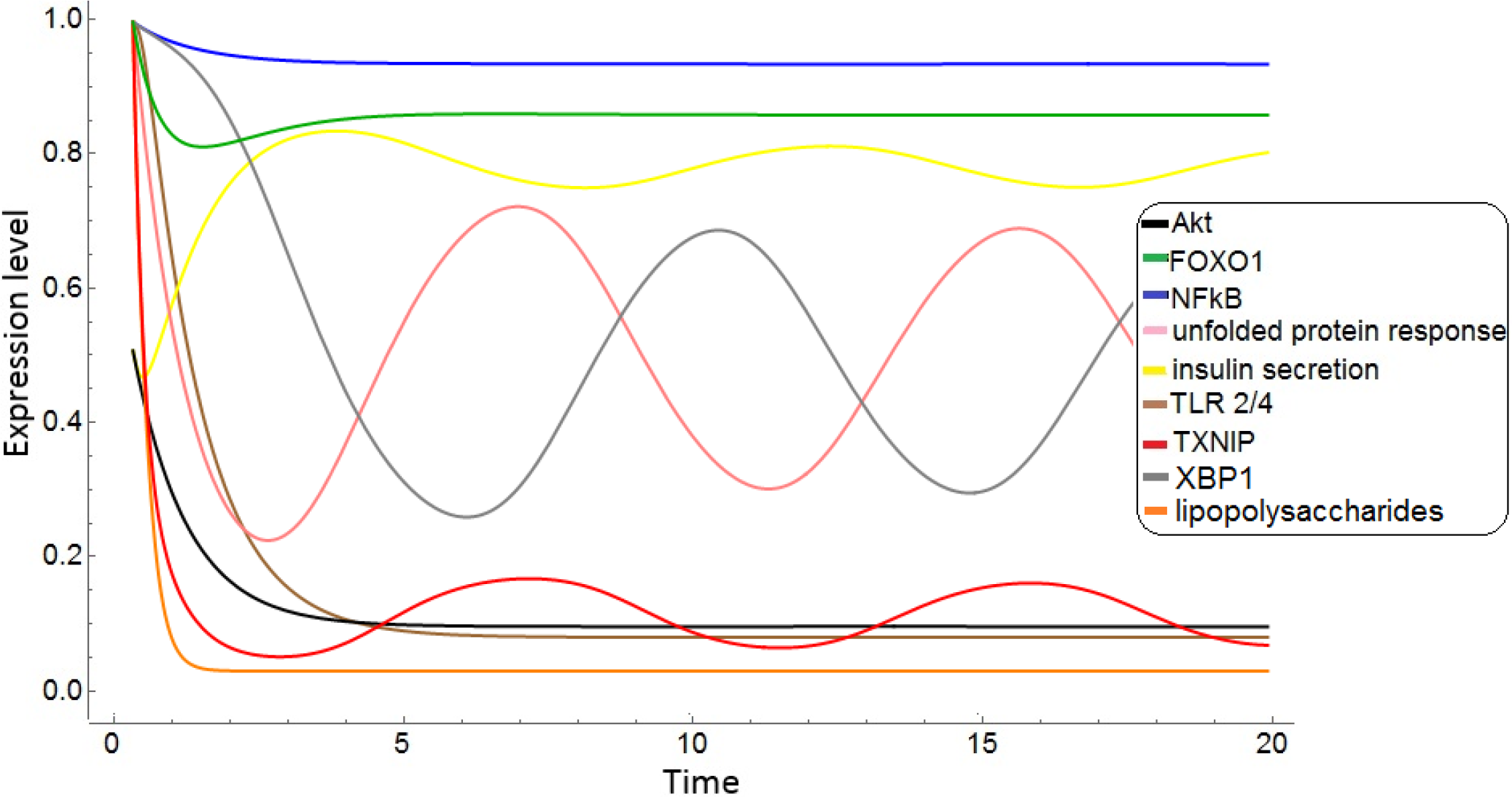
From advanced to improved symptomatology. Dynamic evolution of pancreatic-beta cells expression profiles, from an initial state with high levels of endoplasmic reticulum stress and inflammation, and reduced levels of Akt and insulin secretion, arising from under-expression of TXNIP and lipolysaccharides. The decay rates of all nodes in the beta cells network are *α*_*k*_ = 1, except for the transcription factor TXNIP and LPS, with decay rates *α*_*TXNIP*_ = *α*_*LPS*_ = 5. Insulin secretion recovers from an under-expressed to quasi-normal levels with a mild oscillatory behavior; however, Akt does not recover and remains under-expressed. Inflammation persists, while UPR displays oscillations about a decreased mean level.

As in the transition from health to MS, in this case we also observe expression oscillations of several components of the network. Recurring again to the central core displayed Fig. 2., we notice that inhibition of TXNIP induces the expression of insulin secretion and the decrement of UPR. This decrement inhibits the expression of XBP1 by the alternative pathway (dotted line), promoting in turn an increase in the expression of UPR, leading to a recovery of XBP1, and a consequent cyclic behavior.

## 3 Discussion

We integrated a network of pancreatic-beta cell components (transcription factors, cytokines, hormones, etc.) conducing to early manifestations and progression of type 2 diabetes (T2D). The network comprises connected modules representing insulin signaling and glucose internalization, adenosine triphosphate (ATP) production in glycolisis, inflammation pathways, endoplasmic reticulum (ER) stress, and insulin secretion.

The network regulatory relations, expressed as fuzzy logic propositions, generate a cellular dynamics that portrays the disease progression as a transit between attractors corresponding to health, metabolic syndrome (MS), and T2D. These states would correspond to valleys in the epigenetic landscape, characterized in our approach by expression and inhibition patterns of specific cell components. Going further in the landscape perspective, transitions between valleys may be induced by modulation of the landscape associated to alterations of decay rates of specific network components. This is equivalent to consider alterations of a natural hierarchy of characteristic expression times of the cellular factors involved in MS and T2D, inducing in turn over-expression, under-expression, or exhaustion of key factors involved in signaling pathways. According to the results presented in Figures (3)-(5) for the pancreatic-beta network, the key factors whose under-expression (due to an increased decay rate) lead transitions between alternative disease stages are, besides external lipopolysaccharides (LPS), the internal factors: protein kinase B (AkT), X-box binding protein 1 (XBP1), and thioredoxin interacting protein (TXNIP).

The network analysis reveals that two key elements involved in T2D progression are the transcription factors XBP1 and TXNIP. XBP1 is a major component of the unfolded protein response (UPR), crucial for glucose homeostasis and lipid metabolism. Among other functions, XBP1 regulates ER stress, so that underexpression of this protein leads to increased ER stress and inflammation. Compounds selectively targeting the Inositol-requiring enzyme 1 (IRE1*α*)Äì XBP1 pathway aimed to inhibit XBP1 splicing have emerged as a potential approach for treatment of metabolic diseases^73^. On the other hand, TXNIP regulates metabolic homeostasis through multiple mechanisms. It negatively modulates the activity of thioredoxin, a main controller of the cellular redox balance^74^. TXNIP is upregulated in diabetes, whereas TXNIP deficiency protects against type 1 and T2D by preventing beta cell apoptosis^75^, thus emerging as a novel therapeutic target for the promotion of endogenous-cell mass and insulin production^74, 76^. Experimental studies have provided genetic and pharmacological evidence of the beneficial effects of TXNIP inhibition and its ability to, not only prevent, but also improve overt diabetes^76, 77^. It has been recently documented that calcium channel blockers (i.e., verapamil) inhibit TXNIP expression and beta cell apoptosis^77^. The possibility of ameliorating the symptomatology of the disease through TXNIP inhibition agrees with the theoretical predictions presented above (see Figure 5).

The findings of this study are naturally limited by the pancreatic-beta network design considered here. This network may be complemented by inclusion of supplementary elements that may provide a more detailed description of the cellular mechanisms leading to T2D. In a forthcoming analysis we will investigate the consequences of connecting this network with a regulatory network for hepatocytes on the progression of the disease.

## Conflict of Interest Statement

The authors declare that the research was conducted in the absence of any commercial or financial relationships that could be construed as a potential conflict of interest.

## Author Contributions

M. B., G.C., M.H, and C.V. contributed to the conception of the model. M.B., M. H., and C.V. designed the regulatory network. M.B. constructed the logical propositions, conducted numerical experiments, and performed the analysis of the system dynamics. All authors contributed to the interpretation of the results. M.B. and C.V. wrote the manuscript. All authors reviewed the manuscript.

## Acknowledgments

M. B. is a PhD student from Programa de Doctorado en Ciencias Biomédicas, Universidad Nacional Autónoma de México (UNAM) and received fellowship 379165 from CONACYT. C. V. acknowledges financial support from project CONACYT 180381. All authors acknowledge support from Centro de Ciencias de la Complejidad, UNAM

