## Supplementary Material 1 for "Characterisation of type 2 diabetes progression with regulatory networks"

frontiers

### *Supplementary Material 1*: Characterization of type 2 diabetes progression with regulatory network

#### 1 FORTRAN CODE

```
Program beta cell
! variable statement:
INTEGER n, k, l, ll, i, p, r, s, g, f
INTEGER, DIMENSION (:,:), ALLOCATABLE :: X ! bi-dimensional matrix array
INTEGER, DIMENSION (:,:), ALLOCATABLE :: Y ! one-dimensional vector
INTEGER, DIMENSION (:,:), ALLOCATABLE :: Z ! one-dimensional vector
INTEGER, DIMENSION (:,:), ALLOCATABLE :: W, WW ! one-dimensional vector
INTEGER, DIMENSION (:), ALLOCATABLE :: m, lu, lo
CHARACTER, DIMENSION (:), ALLOCATABLE :: nodes ! constant value

  n = 11 ! number of independent nodes (external and central)
p = 35 ! number of total nodes
r = p-n ! number of dependent nodes

  ALLOCATE (m(n)) ! vector m has n entries
ALLOCATE (nodes(p)) ! vector nodes has p entries

  ! print the total number of configurations of possible initial conditions:
DO i=1,n+1
m(i) = 2^(i − 1)
print*, m(i)
ENDDO

  ! statement of the entries for each array:
ALLOCATE (X(m(n+1),n),Y(m(n+1),p),Z(m(n+1)+1,m(n+1)+1),W(m(n+1)+1,m(n+1)+1),lu(m(n+1)),
lo(m(n+1)),WW(m(n+1)+1,m(n+1)+1))
X = 0
Z = 0
WW = 0
lu = 0
lo = 0
```

! In this section, the program creates the file 'initial conditions.dat' that prints a matrix containing all possible configurations of initial conditions. This matrix is an array of 11 columns and $2^{11}$ rows.

```
open(UNIT=1,FILE ='initial conditions.dat')
DO j=1,n
DO i=1,m(n+1)
l = 0
IF (MOD(i-1,m(j)).EQ.0) THEN
l = (i-1)/m(j)
ENDIF
IF (MOD(l,2).NE.0) THEN
DO k=1,m(j)
X(i+k-1,n+1-j) = 1
ENDDO
ENDIF
ENDDO
ENDDO
```

```
DO i=1,m(n+1)
WRITE(1,*) (X(i,j), j=1,n)
ENDDO
```

! In this section, the program creates the file 'finals.dat', for the set of final stationary conditions associated to every possible set of initial conditions. This array contains 24 columns (r = p - n = 24) and $2^{11}$ rows. Every vector is a result of the following Boolean rules:

```
open(UNIT=2,FILE = 'finals.dat')
DO i=1,m(n+1)
```

Y(i,insulin secretion)=((X(i,insulin) + X(i,extracellular glucose)-X(i,insulin) * X(i,extracellular glucose) + X(i,GLP-1) + X(i,GIP) - X(i,GLP-1) * X(i,GIP)) - (X(i,insulin) + X(i,extracellular glucose) -X(i,insulin) * X(i,extracellular glucose)) * (X(i,GLP-1) + X(i,GIP) - X(i,GLP-1) * X(i,GIP)) + X(i,parasympathetic neuropeptides) + X(i,unfolded protein response) - X(i,parasympathetic neuropeptides) * X(i,unfolded protein response) - ((X(i,insulin) + X(i,extracellular glucose) - X(i,insulin) * X(i,extracellular glucose) + X(i,GLP-1) + X(i,GIP) - X(i,GLP-1) * X(i,GIP)) - (X(i,insulin) + X(i,extracellular glucose) - X(i,insulin) * X(i,extracellular glucose)) * (X(i,GLP-1) + X(i,GIP) - X(i,GLP-1) * X(i,GIP))) * (X(i,parasympathetic neuropeptides) + X(i,unfolded protein response) - X(i,parasympathetic neuropeptides) * X(i,unfolded protein response))) * (1-X(i,ghrelin)) * (1-X(i,sympathetic neuropeptides)) * (1-X(i,unfolded protein response))
Y(i,GLUT2) = X(i,extracellular glucose)
Y(i,intracellular glucose) = X(i,extracellular glucose)
Y(i,NF$\kappa$B) = (X(i,IL1$\beta$/IL6/TNF$\alpha$) + (1-X(i,Akt)) * X(i,extracellular glucose) - X(i,IL1$\beta$/IL6/TNF$\alpha$) * (1-X(i,Akt)) * X(i,extracellular glucose) + X(i,unfolded protein response) + X(i,lipopolysaccharides) - X(i,unfolded protein response) * X(i,lipopolysaccharides)) -(X(i,IL1$\beta$/IL6/TNF$\alpha$) + (1-X(i,Akt))

* X(i,extracellular glucose) - X(i,IL1$\beta$/IL6/TNF$\alpha$) * (1-X(i,Akt)) * X(i,extracellular glucose)) * (X(i,unfolded protein response) + X(i,lipopolysaccharides) - X(i,unfolded protein response) * X(i,lipopolysaccharides))
Y(i,FOXO1) = (1-X(i,Akt)) * X(i,extracellular glucose)
Y(i,AMPK) = 1 - X(i,extracellular glucose)
Y(i,pyruvate) = X(i,extracellular glucose)
Y(i,citric acid cycle) = X(i,extracellular glucose)
Y(i,ATP) = X(i,extracellular glucose)
Y(i,TXNIP) = X(i,unfolded protein response)
Y(i,TLR $2/4$) = X(i,lipopolysaccharides)
Y(i,proinsulin biosynthesis) = X(i,unfolded protein response)
Y(i,IRE1$\alpha$) = X(i,unfolded protein response)
Y(i,ATF6) = X(i,unfolded protein response)
Y(i,PERK) = X(i,unfolded protein response)
Y(i,XBP1) = (X(i,Akt) + X(i,unfolded protein response) - X(i,Akt) * X(i,unfolded protein response)) * (1 - X(i,unfolded protein response))
Y(i,hIRE1$\alpha$) = X(i,unfolded protein response)
Y(i,XBP1s) = X(i,unfolded protein response)
Y(i,eIF2$\alpha$) = X(i,unfolded protein response)
Y(i,ATF4) = X(i,unfolded protein response)
Y(i,chaperone proteins) = X(i,unfolded protein response)
Y(i,JNK) = X(i,unfolded protein response)
Y(i,ER expansion) = X(i,unfolded protein response)
Y(i,apoptosis) = X(i,unfolded protein response)

ENDDO

DO i=1,m(n+1)
WRITE(2,*) (Y(i,j), j=n+1,p)
ENDDO

! In this section a file named 'comparison.dat' is created and this shows the initial conditions that generate the same attractor. This is carried out by comparing all the final conditions in UNIT 2.

open(UNIT=3,FILE= 'comparison.dat')
g = 1
mu = 1
DO i=1,m(n+1)
W(1,i) = i
ENDDO
lu(1) = m(n+1)

```
100 k = mu
l = 0
ll = 0
IF (k.eq.0) THEN
print*, ”error”
ELSE
g = g + 1
DO i=k,k+lu(mu)-1
s=0
DO j=n,p
s= s + ABS(Y(k,j)-Y(W(g-1,i-k+1),j))
ENDDO
IF (s.eq.0) THEN
l = l +1
Z(g-1,l) = W(g-1,i-k+1)
ELSE
ll = ll + 1
W(g,ll) = W(g-1,i-k+1)
ENDIF
ENDDO
ENDIF
mu = W(g,1)
lu(mu) = ll

IF (mu.ne.0) THEN
lo(mu) = l
GOTO 100
ENDIF

DO k= 1, g-2
WRITE(3,*) (Z(k,j), j=1,lo(W(k+1,1)))
WRITE(3,*) ”——————————”

ENDDO

close(1)
close(2)
close(3)

end program beta cell
```
