## Supplemental Data 1 for "Characterisation of type 2 diabetes progression with regulatory networks"

frontiers

### *Supplementary Material 2*: Characterization of type 2 diabetes progression with regulatory network

| Node/Attractor size | 256 | 256 | 128 | 128 | 128 | 128 | 48 | 48 | 15 | 15 |
|---|---|---|---|---|---|---|---|---|---|---|
| **Insulin secretion** | | | | | | | | | | |
| Insulin | - | - | - | - | - | - | - | - | - | - |
| Ghrelin | - | - | - | - | - | - | - | - | - | - |
| Sympathetic neuropeptides | - | - | - | - | - | - | - | - | - | - |
| Parasympathetic neuropeptides | - | - | - | - | - | - | - | - | - | - |
| GLP-1 | - | - | - | - | - | - | - | - | - | - |
| GIP | - | - | - | - | - | - | - | - | - | - |
| Insulin secretion | 0 | 0 | 0 | 0 | 0 | 0 | 0 | 0 | 0 | 1 |
| **Intracellular glucose** | | | | | | | | | | |
| Extracellular glucose | 0 | 0 | 1 | 1 | 1 | 1 | 1 | 1 | 0 | 0 |
| GLUT2 | 0 | 0 | 1 | 1 | 1 | 1 | 1 | 1 | 0 | 0 |
| Intracellular glucose | 0 | 0 | 1 | 1 | 1 | 1 | 1 | 1 | 0 | 0 |
| **ATP production in glycolisis** | | | | | | | | | | |
| Pyruvate | 0 | 0 | 1 | 1 | 1 | 1 | 1 | 1 | 0 | 0 |
| Citric acid cycle | 0 | 0 | 1 | 1 | 1 | 1 | 1 | 1 | 0 | 0 |
| ATP | 0 | 0 | 1 | 1 | 1 | 1 | 1 | 1 | 0 | 0 |
| AMPK | 1 | 1 | 0 | 0 | 0 | 0 | 0 | 0 | 1 | 1 |
| **Inflammation** | | | | | | | | | | |
| IL1$\beta$/IL6/TNF$\alpha$ | - | - | - | - | - | - | - | - | - | - |
| Lipopolysaccharides | 1 | 0 | 0 | 1 | 0 | 0 | 0 | 0 | 0 | 0 |
| TLR 2/4 | 1 | 0 | 0 | 1 | 0 | 0 | 0 | 0 | 0 | 0 |
| FOXO1 | 0 | 0 | 1 | 1 | 0 | 1 | 0 | 0 | 0 | 0 |
| NF$\kappa$B | 1 | 1 | 1 | 1 | 1 | 1 | 1 | 0 | 0 | 0 |
| TXNIP | 1 | 1 | 1 | 1 | 1 | 1 | 0 | 0 | 0 | 0 |
| **Endoplasmic reticulum stress** | | | | | | | | | | |
| Unfolded protein response | - | - | - | - | - | - | - | - | - | - |
| Akt | 0 | 0 | 0 | 0 | 0 | 0 | 0 | 1 | 0 | 0 |
| XBP1 | 0 | 0 | 0 | 0 | 0 | 0 | 1 | 1 | 0 | 0 |
| ATF6 | 1 | 1 | 1 | 1 | 1 | 1 | 0 | 0 | 0 | 0 |
| Chaperone proteins | 1 | 1 | 1 | 1 | 1 | 1 | 0 | 0 | 0 | 0 |
| ER expansion | 1 | 1 | 1 | 1 | 1 | 1 | 0 | 0 | 0 | 0 |
| IRE1$\alpha$ | 1 | 1 | 1 | 1 | 1 | 1 | 0 | 0 | 0 | 0 |
| Proinsulin biosynthesis | 1 | 1 | 0 | 1 | 1 | 1 | 0 | 0 | 0 | 0 |
| PERK | 1 | 1 | 1 | 1 | 1 | 1 | 0 | 0 | 0 | 0 |
| eIF2$\alpha$ | 1 | 1 | 1 | 1 | 1 | 1 | 0 | 0 | 0 | 0 |
| ATF4 | 1 | 1 | 1 | 1 | 1 | 1 | 0 | 0 | 0 | 0 |
| XBP1s | 1 | 1 | 1 | 1 | 1 | 1 | 0 | 0 | 0 | 0 |
| hIRE1$\alpha$ | 1 | 1 | 1 | 1 | 1 | 1 | 0 | 0 | 0 | 0 |
| JNK | 1 | 1 | 1 | 1 | 1 | 1 | 0 | 0 | 0 | 0 |
| Apoptosis | 1 | 1 | 1 | 1 | 1 | 1 | 0 | 0 | 0 | 0 |
| Associated steady state | T2D | T2D | T2D | T2D | T2D | T2D | none | none | none | none |

| Node / Attractor size | 30 | 32 | 32 | 32 | 16 | 16 | 96 | 96 | 96 |
|---|---|---|---|---|---|---|---|---|---|
| **Insulin secretion** | | | | | | | | | |
| Insulin | - | - | - | - | - | - | - | - | - |
| Ghrelin | - | - | - | - | - | - | - | - | - |
| Sympathetic neuropeptides | - | - | - | - | - | - | - | - | - |
| Parasympathetic neuropeptides | - | - | - | - | - | - | - | - | - |
| GLP-1 | - | - | - | - | - | - | - | - | - |
| GIP | - | - | - | - | - | - | - | - | - |
| Insulin secretion | 1 | 1 | 1 | 1 | 1 | 1 | 0 | 0 | 0 |
| **Intracellular glucose** | | | | | | | | | |
| Extracellular glucose | 0 | 1 | 1 | 1 | 1 | 1 | 1 | 1 | 1 |
| GLUT2 | 0 | 1 | 1 | 1 | 1 | 1 | 1 | 1 | 1 |
| Intracellular glucose | 0 | 1 | 1 | 1 | 1 | 1 | 1 | 1 | 1 |
| **ATP production in glycolisis** | | | | | | | | | |
| Pyruvate | 0 | 1 | 1 | 1 | 1 | 1 | 1 | 1 | 1 |
| Citric acid cycle | 0 | 1 | 1 | 1 | 1 | 1 | 1 | 1 | 1 |
| ATP | 0 | 1 | 1 | 1 | 1 | 1 | 1 | 1 | 1 |
| AMPK | 1 | 0 | 0 | 0 | 0 | 0 | 0 | 0 | 0 |
| **Inflammation** | | | | | | | | | |
| IL1$\beta$/IL6/TNF$\alpha$ | - | - | - | - | - | - | - | - | - |
| Lipopolysaccharides | 1 | 1 | 1 | 0 | 0 | 0 | 1 | 1 | 0 |
| TLR 2/4 | 1 | 1 | 1 | 0 | 0 | 0 | 1 | 1 | 0 |
| FOXO1 | 0 | 0 | 1 | 1 | 0 | 0 | 0 | 1 | 1 |
| NF$\kappa$B | 1 | 1 | 1 | 1 | 0 | 1 | 1 | 1 | 1 |
| TXNIP | 0 | 0 | 0 | 0 | 0 | 0 | 0 | 0 | 0 |
| **Endoplasmic reticulum stress** | | | | | | | | | |
| Unfolded protein response | - | - | - | - | - | - | - | - | - |
| Akt | 0 | 0 | 0 | 0 | 1 | 0 | 0 | 0 | 0 |
| XBP1 | 1 | 1 | 0 | 0 | 1 | 1 | 1 | 0 | 0 |
| ATF6 | 0 | 0 | 0 | 0 | 0 | 0 | 0 | 0 | 0 |
| Chaperone proteins | 0 | 0 | 0 | 0 | 0 | 0 | 0 | 0 | 0 |
| ER expansion | 0 | 0 | 0 | 0 | 0 | 0 | 0 | 0 | 0 |
| IRE1$\alpha$ | 0 | 0 | 0 | 0 | 0 | 0 | 0 | 0 | 0 |
| Proinsulin biosynthesis | 0 | 0 | 0 | 0 | 0 | 0 | 0 | 0 | 0 |
| PERK | 0 | 0 | 0 | 0 | 0 | 0 | 0 | 0 | 0 |
| eIF2$\alpha$ | 0 | 0 | 0 | 0 | 0 | 0 | 0 | 0 | 0 |
| ATF4 | 0 | 0 | 0 | 0 | 0 | 0 | 0 | 0 | 0 |
| XBP1s | 0 | 0 | 0 | 0 | 0 | 0 | 0 | 0 | 0 |
| hIRE1$\alpha$ | 0 | 0 | 0 | 0 | 0 | 0 | 0 | 0 | 0 |
| JNK | 0 | 0 | 0 | 0 | 0 | 0 | 0 | 0 | 0 |
| Apoptosis | 0 | 0 | 0 | 0 | 0 | 0 | 0 | 0 | 0 |
| Associated steady state | MS | MS | MS | MS | health | health | none | none | none |

| Node / Attractor size | 15 | 15 | 30 | 49 | 49 | 49 | 98 | 98 | 49 |
|---|---|---|---|---|---|---|---|---|---|
| **Insulin secretion** | | | | | | | | | |
| Insulin | - | - | - | - | - | - | - | - | - |
| Ghrelin | - | - | - | - | - | - | - | - | - |
| Sympathetic neuropeptides | - | - | - | - | - | - | - | - | - |
| Parasympathetic neuropeptides | - | - | - | - | - | - | - | - | - |
| GLP-1 | - | - | - | - | - | - | - | - | - |
| GIP | - | - | - | - | - | - | - | - | - |
| Insulin secretion | 1 | 1 | 1 | 0 | 0 | 0 | 0 | 0 | 0 |
| **Intracellular glucose** | | | | | | | | | |
| Extracellular glucose | 0 | 0 | 0 | 0 | 0 | 0 | 0 | 0 | 0 |
| GLUT2 | 0 | 0 | 0 | 0 | 0 | 0 | 0 | 0 | 0 |
| Intracellular glucose | 0 | 0 | 0 | 0 | 0 | 0 | 0 | 0 | 0 |
| **ATP production in glycolisis** | | | | | | | | | |
| Pyruvate | 0 | 0 | 0 | 0 | 0 | 0 | 0 | 0 | 0 |
| Citric acid cycle | 0 | 0 | 0 | 0 | 0 | 0 | 0 | 0 | 0 |
| ATP | 0 | 0 | 0 | 0 | 0 | 0 | 0 | 0 | 0 |
| AMPK | 1 | 1 | 1 | 1 | 1 | 1 | 1 | 1 | 1 |
| **Inflammation** | | | | | | | | | |
| IL1$\beta$/IL6/TNF$\alpha$ | - | - | - | - | - | - | - | - | - |
| Lipopolysaccharides | 0 | 0 | 1 | 0 | 0 | 0 | 1 | 1 | 0 |
| TLR 2/4 | 0 | 0 | 1 | 0 | 0 | 0 | 1 | 1 | 0 |
| FOXO1 | 0 | 0 | 0 | 0 | 0 | 0 | 0 | 0 | 0 |
| NF$\kappa$B | 1 | 1 | 1 | 0 | 1 | 1 | 1 | 1 | 0 |
| TXNIP | 0 | 1 | 0 | 0 | 0 | 0 | 0 | 0 | 0 |
| **Endoplasmic reticulum stress** | | | | | | | | | |
| Unfolded protein response | - | - | - | - | - | - | - | - | - |
| Akt | 0 | 0 | 0 | 1 | 0 | 0 | 0 | 0 | 0 |
| XBP1 | 0 | 0 | 0 | 1 | 0 | 1 | 0 | 1 | 0 |
| ATF6 | 0 | 0 | 0 | 0 | 0 | 0 | 0 | 0 | 0 |
| Chaperone proteins | 0 | 0 | 0 | 0 | 0 | 0 | 0 | 0 | 0 |
| ER expansion | 0 | 0 | 0 | 0 | 0 | 0 | 0 | 0 | 0 |
| IRE1$\alpha$ | 0 | 0 | 0 | 0 | 0 | 0 | 0 | 0 | 0 |
| Proinsulin biosynthesis | 0 | 0 | 0 | 0 | 0 | 0 | 0 | 0 | 0 |
| PERK | 0 | 0 | 0 | 0 | 0 | 0 | 0 | 0 | 0 |
| eIF2$\alpha$ | 0 | 0 | 0 | 0 | 0 | 0 | 0 | 0 | 0 |
| ATF4 | 0 | 0 | 0 | 0 | 0 | 0 | 0 | 0 | 0 |
| XBP1s | 0 | 0 | 0 | 0 | 0 | 0 | 0 | 0 | 0 |
| hIRE1$\alpha$ | 0 | 0 | 0 | 0 | 0 | 0 | 0 | 0 | 0 |
| JNK | 0 | 0 | 0 | 0 | 0 | 0 | 0 | 0 | 0 |
| Apoptosis | 0 | 0 | 0 | 0 | 0 | 0 | 0 | 0 | 0 |
| Associated steady state | MS | MS | MS | none | none | none | none | none | none |

Table 1. Steady states arising in the Boolean approximation corresponding to health, metabolic syndrome (MS), type 2 diabetes (T2D) or not similar to a known physiological state (denoted as 'none') in beta cells. The table is divided into functional modules corresponding to: insulin secretion, intracellular glucose, ATP production in glycolisis, inflammation and endoplasmic reticulum stress. Nodes that may take alternative values 0 or 1, are denoted as '-'. They represent initial conditions associated to either external inputs or nodes that are considered as central and drive the behavior of the rest through the Boolean interactive rules. Nodes with a defined value represent dependent nodes (extracellular glucose and LPS are independent nodes; however, the interaction rules imply that they must display the explicit values indicated in the table). The header of each column represents the times an attractor is reached. For example, in column 1, the same final state (determined by the value of the dependent nodes) was repeated 256 times.
