## Supplementary Material 3 for "Characterisation of type 2 diabetes progression with regulatory networks"

### *Supplementary Material 3*: Characterization of type 2 diabetes progression with regulatory network

#### 1 FUZZY LOGIC PROPOSITION

The interactions of the beta-pancreatic nodes in the network are characterized by the following set of fuzzy logic propositions. The logical operations **or**, **and**, **not**, are represented by the logical operators $\cup$, $\cap$, $\neg$, respectively.

$$\text{insulin}(t+1) = \text{insulin}(t) \tag{S1}$$

$$\text{Akt}(t+1) = \text{insulin}(t) \cap \neg\text{NF}\kappa\text{B}(t) \tag{S2}$$

$$\begin{aligned}\text{NF}\kappa\text{B}(t)(t+1) &= (\text{IL1}\beta\text{(t)/IL6(t)/TNF}\alpha(t)) \cup \text{TLR } 2/4\text{(t)} \cup \text{TXNIP}(t)\\ &\cup\ \text{FOXO1}(t)\end{aligned} \tag{S3}$$

$$\text{TLR } 2/4(t+1) = \text{Lipopolysaccharides}(t) \tag{S4}$$

$$\text{lipopolysaccharides}(t+1) = \text{Lipopolysaccharides}(t) \tag{S5}$$

$$\text{TXNIP}(t+1) = \text{Unfolded protein response}(t) \tag{S6}$$

$$(\text{IL1}\beta/\text{IL6/TNF}\alpha)(t+1) = (\text{IL1}\beta/\text{IL6/TNF}\alpha)(t) \tag{S7}$$

$$\text{XBP1}(t+1) = \text{Akt}(t) \cup \text{ATF4}(t) \cap \neg\text{ER expansion}(t) \tag{S8}$$

$$\text{ATF4}(t+1) = \text{eIF2}\alpha(t) \tag{S9}$$

$$\text{ER expansion}(t+1) = \text{chaperone proteins}(t) \tag{S10}$$

$$\text{chaperone proteins}(t+1) = \text{XBP1s}(t) \cup \text{ATF6}(t) \tag{S11}$$

$$\text{XBP1s}(t+1) = \text{hIRE1}\alpha(t) \tag{S12}$$

$$\text{unfolded protein response}(t+1) = \text{NF}\kappa\text{B}(t) \cup \text{TXNIP}(t) \cap \neg\text{XBP1}(t) \tag{S13}$$

$$\text{FOXO1}(t+1) = \neg\text{Akt}(t) \cap \neg\text{AMPK}(t) \tag{S14}$$

$$\text{AMPK}(t+1) = \neg\text{ATP}(t) \tag{S15}$$

$$\text{ATP}(t+1) = \text{citric acid cycle}(t) \tag{S16}$$

$$\text{citric acid cycle}(t+1) = \text{pyruvate}(t) \tag{S17}$$

$$\text{pyruvate}(t+1) = \text{intracellular glucose}(t) \tag{S18}$$

$$\text{intracellular glucose}(t+1) = \text{GLUT2}(t) \tag{S19}$$

$$\text{GLUT2}(t+1) = \text{extracellular glucose}(t) \tag{S20}$$

$$
\begin{aligned}
\text{insulin secretion}(t+1) &= \text{insulin}(t) \cup \text{parasympathetic neuropeptides}(t) \\
&\cup\ \text{GLP1}(t) \cup \text{GIP}(t) \cup \text{ATP}(t) \cup \text{proinsulin biosynthesis}(t) \\
&\cap\ \neg\text{ghrelin}(t) \cap \neg\text{sympathetic neuropeptides}(t) \\
&\cap\ \neg\text{TXNIP}(t) && \text{(S21)} \\
\text{proinsulin biosynthesis}(t+1) &= \text{IRE1}\alpha && \text{(S22)} \\
\text{ghrelin}(t+1) &= \text{ghrelin}(t) && \text{(S23)} \\
\text{sympathetic neuropeptides}(t+1) &= \text{sympathetic neuropeptides}(t) && \text{(S24)} \\
\text{parasympathetic neuropeptides}(t+1) &= \text{parasympathetic neuropeptides}(t) && \text{(S25)} \\
\text{GLP-1}(t+1) &= \text{GLP-1}(t) && \text{(S26)} \\
\text{GIP}(t+1) &= \text{GIP}(t) && \text{(S27)} \\
\text{extracellular glucose}(t+1) &= \text{extracellular glucose}(t) && \text{(S28)} \\
\text{IRE1}\alpha(t+1) &= \text{unfolded protein response}(t) && \text{(S29)} \\
\text{hIRE1}\alpha(t+1) &= \text{unfolded protein response}(t) && \text{(S30)} \\
\text{ATF6}(t+1) &= \text{unfolded protein response}(t) && \text{(S31)} \\
\text{PERK}(t+1) &= \text{unfolded protein response}(t) && \text{(S32)} \\
\text{eIF2}\alpha(t+1) &= \text{PERK}(t) && \text{(S33)} \\
\text{JNK}(t+1) &= \text{hIRE1}\alpha(t) && \text{(S34)} \\
\text{apoptosis}(t+1) &= \text{JNK}(t) \cup \text{XBP1s}(t) \cup \text{ATF4}(t) && \text{(S35)}
\end{aligned}
$$
