## Supplementary Material 4 for "Characterisation of type 2 diabetes progression with regulatory networks"

### *Supplementary Material 4*: Characterization of type 2 diabetes progression with regulatory network

#### 1 CONTINUOUS LOGICAL ANALYSIS

Fuzzy logic propositions may be built by replacing ordinary Boolean operations by fuzzy connectors. In this work we consider the following rules for fuzzy logic operations:

$$q \text{ and } p \rightarrow q \cdot p \tag{S1}$$

$$q \text{ or } p \rightarrow q + p - q \cdot p \tag{S2}$$

$$\text{not } p \rightarrow 1 - p \tag{S3}$$

For example, the proposition $q$ or $p$ and (not $z$) $\rightarrow (q + p - q \cdot p) \cdot (1 - z)$. A straightforward consequence of the former rules is that, since the no-contradiction principle is not satisfied in fuzzy logic, a proposition $q$ and its negation $1 - q$ may be simultaneously true. In that case, $q = 1 - q$, with solution $q = 1/2 \equiv q^{thr}$. Thus, $q^{thr}$ may be interpreted as a threshold between falsity and truth, which we employ in the following.

Consider now a regulatory network consisting of $n$ interacting nodes with expression levels at a given time $t$ represented by $q_k(t)$, with $k = 1, ..., n$. The state of node $k$ is regulated by its interaction with the rest of the network nodes, determined by a fuzzy proposition $w_k(q_1(t), ..., q_n(t))$. This proposition may be either inferred from experimental observations, or suggested by inner consistency requirements. Within this approach, the expression degree (truth level) of $w_k$ is determined by its characteristic (or membership) function $\mu(w_k)$, which may be evaluated by considering a generalization of the central argument involved in logistic regression analysis. Following similar lines, we introduce the odds $\mu(w_k)/[1 - \mu(w_k)]$, that is, the ratio of possibilities that $\mu(w_k)$ is either true or false, (expressed or non-expressed). We assume that the natural logarithm of the odds is proportional to the condition that $w_k$ is greater or equal than a threshold level $w^{thr}$:

$$\ln\left[\mu(w_k)/(1 - \mu(w_k))\right] = b(w_k - w^{thr}), \tag{S4}$$

By solving this latter expression for $\mu(w_k)$, we obtain a logistic expression for the characteristic function:

$$\mu(w_k) = \frac{1}{1 + \exp\left[-b\left(w_k(q_1, ..., q_n) - w^{thr}\right)\right]}. \tag{S5}$$

Here, $b \geq 1$ is a saturation parameter for the progression pace of $\mu$ from unexpressed to expressed, gradual for small $b$, steep for large $b$. In the case $b \gg 1$, the characteristic function is equivalent to a differentiable step function:

$$\mu[w_k] \rightarrow \Theta[w_k - w^{thr}] = \begin{cases} 1 & \text{if } w_k > w^{thr}; \\ 1/2 & \text{if } w_k = w^{thr}; \\ 0 & \text{if } w_k < w^{thr}. \end{cases}$$

In this work we supposed that $b = 5$, and $w^{thr} = 1/2$.

The dynamics of the fuzzy network is described by a set of ordinary differential equations for the rate of change of the expression level of the network components. For each node, this rate is given by

$$\frac{dq_k}{dt} = \mu(w_k) - \alpha_k q_k, \tag{S6}$$

where $\alpha_k$ is the decay rate of component $k$ of the network, so that in absence of a regulatory interaction, its expression level suffers an exponential time-decay at a rate $\alpha_k = 1/\tau_k$, where $\tau_k$ is a characteristic expression time. The equilibrium states of the system may be obtained from the steady-state condition $dq_k/dt = 0$, leading to the stationary expression $q_k^s = \mu(w_k^s)/\alpha_k$. We observe that the case $\alpha_k = 1$ for every node, and $b \gg 1$, corresponds (by construction) to the Boolean limit described by a mapping with synchronous updating ($\tau_k = 1$ for every node), since the attractors are given by sets of totally expressed or unexpressed variables, for example, $\{1, 0, 1, ..., \}$. On the other hand, in the more general case $\tau_k$ may be different for every node, and the set $\{\alpha_k\}$ induces a hierarchy of expression times of the regulatory network. This hierarchy induces in turn a modulation of the network expression levels: since the maximal value of $\mu = 1$, when $\alpha_k > 1$, a node will be under-expressed, with $q_k^s < 1$; in particular, for $\alpha_k \gg 1$, a node expression is completely inhibited: $q_k^s \to 0$.

According to the former analysis the expression patterns of the network components may be altered by modifications of their characteristic decay rates. This is equivalent to modulate the effective epigenetic landscape underlying the regulatory network. In this work, this mechanism may induce transitions between attractors associated with different stages involved in T2D progression: health, MS, and T2D manifestations. Thus, in order to study the transitions between these latter states, we first determined the pathways leading from the possible initial configurations of the network to the final attractors set.
