## Supplementary Material 5 for "Characterisation of type 2 diabetes progression with regulatory networks"

### *Supplementary Material 5*: Characterization of type 2 diabetes progression with regulatory network

#### 1 WOLFRAM MATHEMATICA CODE

(*Note: text between the signs (* *) represent comments in the Wolfram Mathematica enviroment.*)

(*As we have mentioned in the main text, we did a dynamical analysis to study the transitions between steady states related to health, metabolic syndrome (MS) and type 2 diabetes (T2D). We used Wolfram Mathematica computing system to solve the ordinary differential equations that describe the expression pattern of every state depicted in Table 1 and the transitions between them. The equation that describe each node evolution is given by the equation 8 (main text):*)

(*

$$\frac{dq_k}{dt} = \mu(w_k) - \alpha_k q_k, \tag{S1}$$

$$\mu(w_k) = \frac{1}{1 + \exp\left[-b\left(w_k(q_1, ..., q_n) - w^{thr}\right)\right]}. \tag{S2}$$

$\mu(w_k)$ is a characteristic function with a logistic behavior, representing the activity level induced by the action of the fuzzy logic proposition $w_k$ acting on the $k$th node. $w^{thr}$ is a threshold value, $w^{thr} = 1/2$. $b$ is a variation rate for the change of the proposition $\mu(w_k)$ from an unexpressed to a totally expressed state. $q_k$ is the expression level of the component $k$.*)

(*The interactions of the beta-pancreatic nodes in the network are characterized by the following set of fuzzy logic propositions.*)

##### 1.1 Fuzzy logic propositions

(*These are the same propositions mentioned in Supplementary material 3, but here, the logical operators ∪, ∩, ¬ were expressed as*)

(*

$$q \cap p \rightarrow q \cdot p \tag{S3}$$

$$q \cup p \rightarrow q + p - q \cdot p \tag{S4}$$

$$\neg p \rightarrow 1 - p \tag{S5}$$

*)

$$\begin{aligned}
w\text{insulin}(t+1) &= \text{insulin}(t) && \text{(S6)}\\
w\text{Akt}(t+1) &= \text{insulin}(t)(1-\text{NF}\kappa\text{B}(t)) && \text{(S7)}\\
w\text{NF}\kappa\text{B}(t)(t+1) &= (\text{IL1}\beta\text{(t)/IL6(t)/TNF}\alpha(t))+\text{FOXO1}(t)\\
&- (\text{IL1}\beta\text{(t)/IL6(t)/TNF}\alpha(t))\text{FOXO1(t)+TXNIP(t)+TLR}2/4(t)\\
&- \text{TXNIP}(t)\text{TLR}2/4(t)-((\text{IL1}\beta\text{(t)/IL6(t)/TNF}\alpha(t))+\text{FOXO1}(t)\\
&- -(\text{IL1}\beta\text{(t)/IL6(t)/TNF}\alpha(t))\text{FOXO1}(t))(\text{TXNIP}(t)+\text{TLR}2/4\\
&- \text{TXNIP}(t)\text{TLR}2/4) && \text{(S8)}\\
w\text{TLR } 2/4(t+1) &= \text{Lipopolysaccharides}(t) && \text{(S9)}\\
w\text{lipopolysaccharides}(t+1) &= \text{Lipopolysaccharides}(t) && \text{(S10)}\\
w\text{TXNIP}(t+1) &= \text{Unfolded protein response}(t) && \text{(S11)}\\
w(\text{IL1}\beta\text{/IL6/TNF}\alpha)(t+1) &= (\text{IL1}\beta\text{/IL6/TNF}\alpha)(t) && \text{(S12)}\\
w\text{XBP1}(t+1) &= (\text{Akt}(t)+\text{ATF4}(t)\text{-Akt}(t)\text{ATF4}(t))(1-\text{ER expansion}(t)) && \text{(S13)}\\
w\text{ATF4}(t+1) &= \text{eIF2}\alpha(t) && \text{(S14)}\\
w\text{ER expansion}(t+1) &= \text{chaperone proteins}(t) && \text{(S15)}\\
w\text{chaperone proteins}(t+1) &= \text{XBP1s}(t)+\text{ATF6}(t)\text{-XBP1s}(t)\text{ATF6}(t) && \text{(S16)}\\
w\text{XBP1s}(t+1) &= \text{hIRE1}\alpha(t) && \text{(S17)}\\
w\text{unfolded protein response}(t+1) &= (\text{NF}\kappa\text{B}(t)+\text{TXNIP}(t)\text{-NF}\kappa\text{B}(t)\text{TXNIP}(t))(1-\text{XBP1(t)}) && \text{(S18)}\\
w\text{FOXO1}(t+1) &= (1-\text{Akt}(t))(1-\text{AMPK}(t)) && \text{(S19)}\\
w\text{AMPK}(t+1) &= (1-\text{ATP}(t)) && \text{(S20)}\\
w\text{ATP}(t+1) &= \text{citric acid cycle}(t) && \text{(S21)}\\
w\text{citric acid cycle}(t+1) &= \text{pyruvate}(t) && \text{(S22)}\\
w\text{pyruvate}(t+1) &= \text{intracellular glucose}(t) && \text{(S23)}\\
w\text{intracellular glucose}(t+1) &= \text{GLUT2}(t) && \text{(S24)}\\
w\text{GLUT2}(t+1) &= \text{extracellular glucose}(t) && \text{(S25)}
\end{aligned}$$

$$\begin{aligned}
w\text{insulin secretion}(t+1) &= (\text{insulin}(t)+\text{ATP}(t)-\text{insulin}(t)\text{ATP}(t)+\text{GIP}(t)+\text{GLP1}(t) \\
&- \text{GIP}(t)\text{GLP1}(t))-(\text{insulin}(t)+\text{ATP}(t)-\text{insulin}(t)\text{ATP}(t))(\text{GIP}(t) \\
&+ \text{GLP1}(t)-\text{GIP}(t)+\text{GLP1}(t))+\text{parasympathetic neuropeptides}(t) \\
&+ \text{proinsulin biosynthesis}(t) \\
&- \text{parasympathetic neuropeptides}(t)\text{proinsulin biosynthesis}(t) \\
&- ((\text{insulin}(t)+\text{ATP}(t)-\text{insulin}(t)\text{ATP}(t)+\text{GIP}(t) \\
&+ \text{GLP1}(t)-\text{GIP}(t)\text{GLP1}(t)) \\
&- (\text{insulin}(t)+\text{ATP}(t)-\text{insulin}(t)\text{ATP}(t))(\text{GIP}(t) \\
&+ \text{GLP1}(t)-\text{GIP}(t)\text{GLP1}(t)))(\text{parasympathetic neuropeptides}(t) \\
&+ \text{proinsulin biosynthesis}(t) \\
&- \text{parasympathetic neuropeptides}(t)\text{proinsulin biosynthesis}(t)) \\
&* (1-\text{ghrelin}(t))(1-\text{sympathetic neuropeptides}(t)) \\
&* (1-\text{TXNIP}(t)) \qquad \text{(S26)}
\end{aligned}$$

$$w\text{proinsulin biosynthesis}(t+1) = \text{IRE1}\alpha \qquad \text{(S27)}$$

$$w\text{ghrelin}(t+1) = \text{ghrelin}(t) \qquad \text{(S28)}$$

$$w\text{sympathetic neuropeptides}(t+1) = \text{sympathetic neuropeptides}(t) \qquad \text{(S29)}$$

$$w\text{parasympathetic neuropeptides}(t+1) = \text{parasympathetic neuropeptides}(t) \qquad \text{(S30)}$$

$$w\text{GLP-1}(t+1) = \text{GLP-1}(t) \qquad \text{(S31)}$$

$$w\text{GIP}(t+1) = \text{GIP}(t) \qquad \text{(S32)}$$

$$w\text{extracellular glucose}(t+1) = \text{extracellular glucose}(t) \qquad \text{(S33)}$$

$$w\text{IRE1}\alpha(t+1) = \text{unfolded protein response}(t) \qquad \text{(S34)}$$

$$w\text{hIRE1}\alpha(t+1) = \text{unfolded protein response}(t) \qquad \text{(S35)}$$

$$w\text{ATF6}(t+1) = \text{unfolded protein response}(t) \qquad \text{(S36)}$$

$$w\text{PERK}(t+1) = \text{unfolded protein response}(t) \qquad \text{(S37)}$$

$$w\text{eIF2}\alpha(t+1) = \text{PERK}(t) \qquad \text{(S38)}$$

$$w\text{JNK}(t+1) = \text{hIRE1}\alpha(t) \qquad \text{(S39)}$$

$$\begin{aligned}
w\text{apoptosis}(t+1) &= \text{JNK}(t)+\text{XBP1s}(t)-\text{JNK}(t)\text{XBP1s}(t)+\text{ATF4}(t) \\
&- (\text{JNK}(t)+\text{XBP1s}(t)-\text{JNK}(t)\text{XBP1s}(t))\text{ATF4}(t) \qquad \text{(S40)}
\end{aligned}$$

#### 1.2 Definitions of decay rates and saturation parameter of fuzzy logic propositions

b $= 5$;
(* saturation parameter b is a variation rate for the change of the fuzzy logic proposition from an unexpressed to a totally expressed state. (Section 2.3 Continuous logical analysis) *).
$\alpha_{insulin}$ = 1;
(* $\alpha_k$=Decay rate of component $k$ of the network). It was initially assumed that $\alpha_k = 1$ for every node. Then, we modified every decay rate from its initial value $\alpha_k = 1$ to a maximal value $\alpha_k = 5$. The next

decay rates show the situation where we detected a transition from health to transient MS, and final manifest T2D (Figure 4).*).
$\alpha_{Akt}$ = 1;
$\alpha_{NFkB}$ = 1;
$\alpha_{TLR2/4}$ = 1.;
$\alpha_{lipopolysaccharides}$ = 1;
$\alpha_{TXNIP}$ = 1;
$\alpha$IL1$\beta$/IL6/TNF$\alpha$ = 1;
$\alpha_{XBP1}$ = 3;
$\alpha_{ATF4}$ = 1;
$\alpha_{ERexpansion}$ = 1;
$\alpha_{chaperoneproteins}$ = 1;
$\alpha_{XBP1s}$ = 1;
$\alpha_{unfoldedproteinresponse}$ = 1;
$\alpha_{FOXO1}$ = 1;
$\alpha_{AMPK}$ = 1;
$\alpha_{ATP}$ = 1;
$\alpha_{citricacidcycle}$= 1;
$\alpha_{pyruvate}$ = 1;
$\alpha_{intracellularglucose}$ = 1;
$\alpha_{GLUT2}$ = 1;
$\alpha_{insulinsecretion}$ = 1;
$\alpha_{proinsulinbiosynthesis}$ = 1;
$\alpha_{ghrelin}$ = 1;
$\alpha_{sympatheticneuropeptides}$ = 1;
$\alpha_{parasympatheticneuropeptides}$ = 1;
$\alpha_{GLP1}$ = 1;
$\alpha_{GIP}$ = 1;
$\alpha_{extracellularglucose}$ = 1;
$\alpha_{IRE1}$ = 1;
$\alpha_{hIRE1}$ = 1;
$\alpha_{ATF6}$ = 1;
$\alpha_{PERK}$ = 1;
$\alpha_{eIF2}$ = 1;
$\alpha_{JNK}$ = 1;
$\alpha_{apoptosis}$ = 1;

##### 1.3 Initial conditions of the network

(*Each attractor in Table 1 was in this step considered as a set of initial conditions of every node. The next initial conditions are an example of the case where the health state was analyzed. *)

insulin0 = 1;
Akt0 = 1;
NFkB0 = 0;
TLR $2/40$ = 0;
lipopolysaccharides0= 0;

TXNIP0 = 0;
IL1$\beta$/IL6/TNF$\alpha$0 = 0;
XBP10 = 0;
ATF40 = 1;
ER expansion0 = 0;
chaperone proteins0 = 0;
XBP1s0 = 0;
unfolded protein response0 = 0;
FOXO10 = 0;
AMPK0 = 1;
ATP0 = 1;
citric acid cycle0 = 1;
pyruvate0 = 1;
intracellular glucose0 = 1;
GLUT20 = 1;
insulin secretion0 = 1;
proinsulin biosynthesis0 = 1;
ghrelin0 = 0;
sympathetic neuropeptides0 = 0;
parasympathetic neuropeptides0 = 1;
GLP10 = 1;
GIP0 = 1;
extracellular glucose0 = 1;
IRE10 = 1;
hIRE10 = 0;
ATF60 = 0;
PERK0 = 0;
eIF20 = 0;
JNK0 = 0;
apoptosis0 = 0;

#### 1.4 Solution of ordinary differential equations

(*The instruction 'NDSolve' solves a differential equation numerically, using the initial conditions previously defined. Then, we showed the solutions in form of plots (Figures 3 to 5, main text). The equation 8 of the main text was solved for every node of the network.*)

netw=NDSolve[insulin'[t] == 1/(1 + Exp[-b (winsulin[t] - .5)]) - $\alpha_{insulin}$ insulin[t],
Akt'[t] == 1/(1 + Exp[-b (wAkt[t] - .5)]) - $\alpha_{Akt}$ Akt[t],
NFkB'[t] == 1/(1 + Exp[-b (wNFkB[t] - .5)]) - $\alpha_{NFkB}$ NFkB[t],
TLR $2/4$'[t] == 1/(1 + Exp[-b (wTLR $2/4$[t] - .5)]) - $\alpha_{TLR2/4}$ TLR $2/4$[t],
lipopolysaccharides'[t] == 1/(1 + Exp[-b (wlipopolysaccharides[t] - .5)]) - $\alpha_{lipopolysaccharides}$ lipopolysaccharides[t],
TXNIP'[t] == 1/(1 + Exp[-b (wTXNIP[t] - .5)]) - $\alpha_{TXNIP}$ TXNIP[t],
IL1$\beta$/IL6/TNF$\alpha$'[t] == 1/(1 + Exp[-b (wIL1$\beta$/IL6/TNF$\alpha$[t] - .5)]) - $\alpha$IL1$\beta$/IL6/TNF$\alpha$ IL1$\beta$/IL6/TNF$\alpha$[t],
XBP1'[t] == 1/(1 + Exp[-b (wXBP1[t] - .5)]) - $\alpha_{XBP1}$ XBP1[t],
ATF4'[t] == 1/(1 + Exp[-b (wATF4[t] - .5)]) - $\alpha_{ATF4}$ ATF4[t],

ER expansion'[t] == 1/(1 + Exp[-b (wER expansion[t] - .5)]) - $\alpha_{ERexpansion}$ ER expansion[t],
chaperoneproteins'[t] == 1/(1 + Exp[-b (wchaperone proteins[t] - .5)]) - $\alpha_{chaperoneproteins}$ chaperone proteins[t],
XBP1s'[t] == 1/(1 + Exp[-b (wXBP1s[t] - .5)]) - $\alpha_{XBP1s}$ XBP1s[t],
unfolded protein response'[t] == 1/(1 + Exp[-b (wunfolded protein response[t] - .5)]) - $\alpha_{unfoldedproteinresponse}$ unfolded protein response[t],
FOXO1'[t] == 1/(1 + Exp[-b (wFOXO1[t] - .5)]) - $\alpha_{FOXO1}$ FOXO1[t],
AMPK'[t] == 1/(1 + Exp[-b (wAMPK[t] - .5)]) - $\alpha_{AMPK}$ AMPK[t],
ATP'[t] == 1/(1 + Exp[-b (wATP[t] - .5)]) - $\alpha_{ATP}$ ATP[t],
citric acid cycle'[t] == 1/(1 + Exp[-b (wcitric acid cycle[t] - .5)]) - $\alpha_{citricacidcycle}$ citric acid cycle[t],
pyruvate'[t] == 1/(1 + Exp[-b (wpyruvate[t] - .5)]) - $\alpha_{pyruvate}$ pyruvate[t],
intracellular glucose'[t] == 1/(1 + Exp[-b (wintracellular glucose[t] - .5)]) - $\alpha_{intracellularglucose}$ intracellular glucose[t],
GLUT2'[t] == 1/(1 + Exp[-b (wGLUT2[t] - .5)]) - $\alpha_{GLUT2}$ GLUT2[t],
insulin secretion'[t] == 1/(1 + Exp[-b (winsulin secretion[t] - .5)]) - $\alpha_{insulinsecretion}$ insulin secretion[t],
proinsulin biosynthesis'[t] == 1/(1 + Exp[-b (wproinsulin biosynthesis[t] - .5)]) - $\alpha_{proinsulinbiosynthesis}$ proinsulin biosynthesis[t],
ghrelin'[t] == 1/(1 + Exp[-b (wghrelin[t] - .5)]) - $\alpha_{ghrelin}$ ghrelin[t],
sympathetic neuropeptides'[t] == 1/(1 + Exp[-b (wsympathetic neuropeptides[t] - .5)]) - $\alpha_{sympatheticneuropeptides}$ sympathetic neuropeptides[t],
parasympathetic neuropeptides'[t] == 1/(1 + Exp[-b (wparasympathetic neuropeptides[t] - .5)]) - $\alpha_{parasympatheticneuropeptides}$ parasympathetic neuropeptides[t],
GLP1'[t] == 1/(1 + Exp[-b (wGLP1[t] - .5)]) - $\alpha_{GLP1}$ GLP1[t],
GIP'[t] == 1/(1 + Exp[-b (wGIP[t] - .5)]) - $\alpha_{GIP}$ GIP[t],
extracellular glucose'[t] == 1/(1 + Exp[-b (wextracellular glucose[t] - .5)]) - $\alpha_{extracellularglucose}$ extracellular glucose[t],
IRE1'[t] == 1/(1 + Exp[-b (wIRE1[t] - .5)]) - $\alpha_{IRE1}$ IRE1[t],
hIRE1'[t] == 1/(1 + Exp[-b (whIRE1[t] - .5)]) - dhIRE1 $\alpha_{hIRE1}$[t],
ATF6'[t] == 1/(1 + Exp[-b (wATF6[t] - .5)]) - $\alpha_{ATF6}$ ATF6[t],
PERK'[t] == 1/(1 + Exp[-b (wPERK[t] - .5)]) - $\alpha_{PERK}$ PERK[t],
eIF2'[t] == 1/(1 + Exp[-b (weIF2[t] - .5)]) - $\alpha_{eIF2}$ eIF2[t],
JNK'[t] == 1/(1 + Exp[-b (wJNK[t] - .5)]) - $\alpha_{JNK}$ JNK[t],
apoptosis'[t] == 1/(1 + Exp[-b (wapoptosis[t] - .5)]) - $\alpha_{apoptosis}$ apoptosis[t],

insulin[0] == insulin0, Akt[0] == Akt0, NFkB[0] == NFkB0, TLR$2/4$[0] == TLR$2/4$0, lipopolysaccharides[0] == lipopolysaccharides0, TXNIP[0] == TXNIP0, IL1$\beta$/IL6/TNF$\alpha$[0] == IL1$\beta$/IL6/TNF$\alpha$0, XBP1[0] == XBP10, ATF4[0] == ATF40, ER expansion[0] == ER expansion0, chaperone proteins[0] == chaperone proteins0, XBP1s[0] == XBP1s0, unfolded protein response[0] == unfolded protein response0, FOXO1[0] == FOXO10, AMPK[0] == AMPK0, ATP[0] == ATP0, citric acid cycle[0] == citric acid cycle0, pyruvate[0] == pyruvate0, intracellular glucose[0] == intracellular glucose0, GLUT2[0] == GLUT20, insulin secretion[0] == insulin secretion0, proinsulin biosynthesis[0] == proinsulin biosynthesis0, ghrelin[0] == ghrelin0, sympathetic neuropeptides[0] == sympathetic neuropeptides0, parasympathetic neuropeptides[0] == parasympathetic neuropeptides0, GLP1[0] == GLP10, GIP[0] == GIP0, extracellular glucose[0] == extracellular glucose0, IRE1[0] == IRE10, hIRE1[0] == hIRE10, ATF6[0] == ATF60, PERK[0] == PERK0, eIF2[0] == eIF20, JNK[0] == JNK0, apoptosis[0] == apoptosis0, insulin[t], Akt[t], NFkB[t], TLR2/4[t], lipopolysaccharides[t], TXNIP[t], IL1$\beta$/IL6/TNF$\alpha$[t], XBP1[t], ATF4[t], ER expansion[t], chaperone proteins[t], XBP1s[t], unfolded protein response[t], FOXO1[t], AMPK[t], ATP[t], citric acid cycle[t], pyruvate[t], intracellular glucose[t], GLUT2[t], insulin secretion[t], proinsulin biosynthesis[t], ghrelin[t], sympathetic neuropeptides[t], parasympathetic neuropeptides[t], GLP1[t], GIP[t], extracellular glucose[t], IRE1[t], hIRE1[t], ATF6[t], PERK[t], eIF2[t], JNK[t], apoptosis[t], t, 0, 20.]

health=Plot[Evaluate[Akt[t], FOXO1[t], NFkB[t], unfolded protein response[t], insulin secretion[t], TLR2/4[t], TXNIP[t], XBP1[t], apoptosis[t] /.netw], t, 0, 20, PlotRange All, FrameLabel , , time, ”Expression level”, PlotStyle Thick, Black, Thick, Darker[Green], Thick, Blue, Thick, Pink, Thick, Yellow, Thick, Brown, Thick, Red, Thick, Gray, Thick, Purple, Frame True, PlotLabel ”Health to transient MS, and final T2D”]

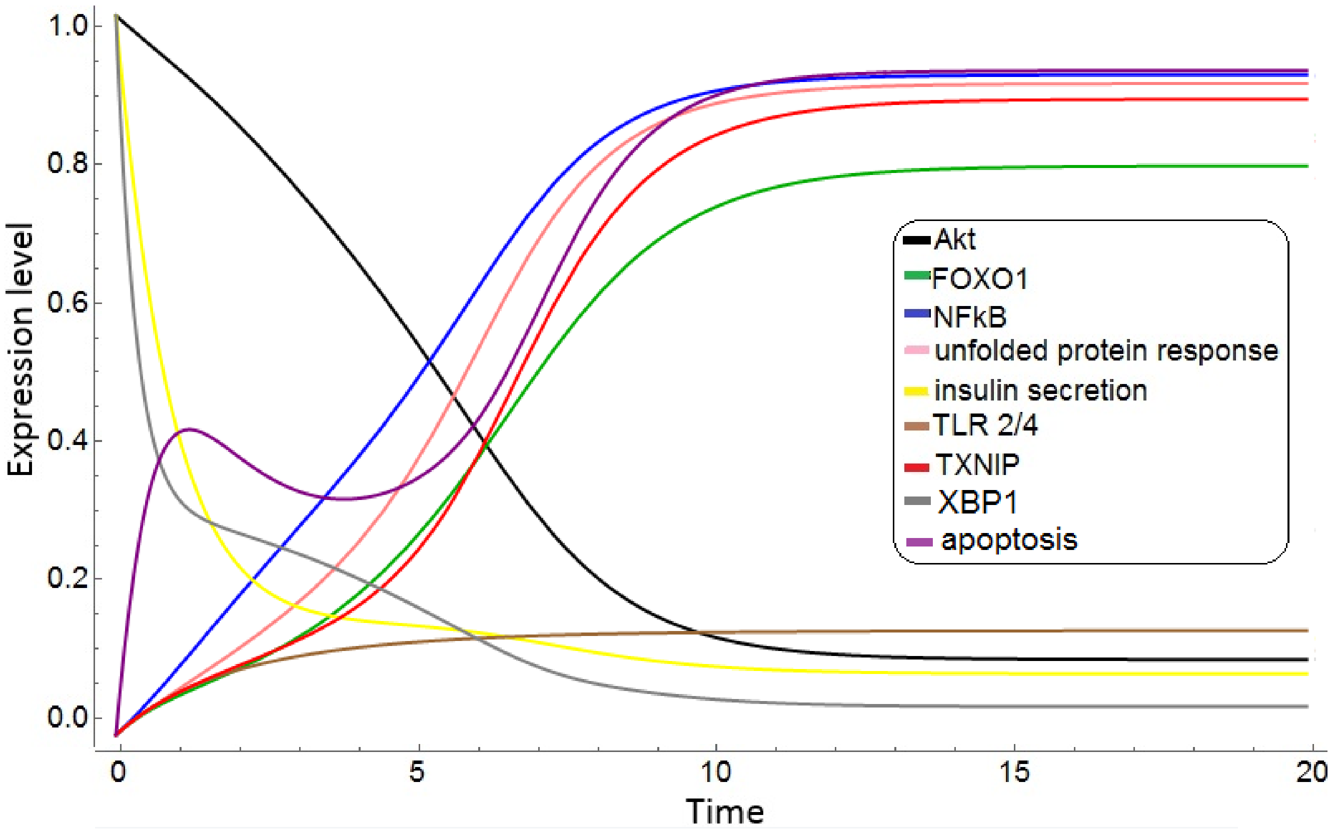


**Figure S1.** From health to transient metabolic syndrome, and subsequent type 2 diabetes. This figure is a result of solving the equation systems with the specific initial conditions and decay rates aforementioned (Figure 4 main text).
